# Transcriptomic analysis reveals niche gene expression effects of beta-hydroxybutyrate in primary myotubes

**DOI:** 10.1101/2021.01.19.427259

**Authors:** Philip M. M. Ruppert, Guido J. E. J. Hooiveld, Roland W. J. Hangelbroek, Anja Zeigerer, Sander Kersten

**Affiliations:** Nutrition, Metabolism and Genomics Group, Division of Human Nutrition and Health, Wageningen University, Wageningen, The Netherlands; Euretos B.V., Utrecht, The Netherlands; Institute for Diabetes and Cancer, Helmholtz Center Munich, 85764 Neuherberg, Germany, Joint Heidelberg-IDC Translational Diabetes Program, Inner Medicine 1, Heidelberg University Hospital, Heidelberg, Germany; German Center for Diabetes Research (DZD), 85764 Neuherberg, Germany

**Author notes:** To whom correspondence should be addressed: Sander Kersten, PhD, Division of Human Nutrition and Health, Wageningen University, Stippeneng 4, 6708 WE Wageningen, The Netherlands.

## Abstract

Various forms of fasting, including time-restricted feeding, alternate day fasting, and periodic fasting have shown promise in clinical and pre-clinical studies to normalize body weight, improve metabolic health, and protect against disease. Recent studies suggest that β-hydroxybutyrate (βOHB), a characteristic ketone body of the fasted metabolic state, acts as a potential signaling molecule mediating the beneficial effects of the various forms of fasting, potentially by acting as a histone deacetylase inhibitor. In the first part we investigated whether βOHB, in comparison to the well-established histone deacetylase inhibitor butyrate, influences cellular differentiation *in vitro*. In C2C12 myotubes, 3T3-L1 adipocytes, and THP-1 monocytes, millimolar concentrations of βOHB did not alter differentiation, as determined by gene expression and histological assessment, whereas equimolar concentrations of butyrate potently impaired differentiation in all cell types. RNA-sequencing revealed that unlike butyrate, βOHB minimally impacted gene expression in adipocytes, macrophages, and hepatocytes. However, in myocytes, βOHB upregulated genes involved in the TCA cycle and oxidative phosphorylation, while downregulating genes belonging to cytokine and chemokine signal transduction. Overall, our data do not support the notion that βOHB serves as a powerful signaling molecule regulating gene expression in adipocytes, macrophages and hepatocytes, but suggest that βOHB may act as a niche signaling molecule in muscle.

## 1. INTRODUCTION

Prevalence rates for obesity are spiraling out of control in many communities across the world. Inasmuch as obesity is a major risk factor for many chronic diseases, including type 2 diabetes, cardiovascular disease, and certain types of cancer [1], effective remedies to slow down the growth of obesity are direly needed. A common strategy that effectively promotes weight loss, at least in the short term, is caloric restriction, leading to an improvement in the cardiometabolic risk profile. One of the more popular forms of caloric restriction is time-restricted feeding, in which the normal abstinence of food consumption during the night is partly extended into the daytime [2]. Other forms of caloric restriction include alternate day fasting, periodic fasting (e.g. 5:2) as well as fasting mimicking diets [2]. In animal models, these dietary interventions increase median life-span, reduce body weight, mitigate inflammation, improve glucose homeostasis and insulin sensitivity, and delay the onset of diabetes, cardiovascular and neurological disease, as well as cancer. Similarly, human studies reported weight loss, reduced HbA1c and glucose levels, improved insulin sensitivity and blood lipid parameters, as well as lower blood pressure [2]–[7].

Interestingly, it has been suggested that intermittent fasting may confer cardiometabolic health benefits independently of caloric restriction and concomitant weight loss [7], [8]. A number of mechanisms have been invoked in explaining the possible health benefits of the various forms of fasting, including lower plasma insulin levels and higher levels of ketone bodies. Ketonemia is a characteristic feature of the fasted metabolic state. During the feeding-fasting transition, the body switches from glucose as a primary fuel source to the oxidation of fatty acids. In the liver, the high rates of fatty acid oxidation are accompanied by the synthesis of ketone bodies, which, as fasting progresses, become the dominant fuel for the brain [9]. The two main ketone bodies are β-hydroxybutyrate (βOHB) and acetoacetate (AcAc) and compounds serve as sensitive biomarkers for the fasted state, increasing in combined concentration from less than 0.1 mM in the fed state to 1 mM after 24h or 5-7 mM when fasting for about a week [9]–[11].

In addition to serving as fuel in tissues such as the brain, heart, and skeletal muscle, recent research has unveiled that βOHB may also serve as a direct signaling molecule. By activating specific signaling pathways, βOHB may not only have an important regulatory role in the metabolic response to fasting but may also potentially mediate some of the beneficial health effects of fasting [2], [12]–[23]. Evidence has been presented that βOHB may regulate gene expression via epigenetic mechanisms. Shimazu *et al*. linked βOHB-mediated HDAC inhibition to protection against oxidative stress in kidney via the up-regulation of *FOXO3a, Catalase* and *MnSOD* [24]. While subsequent studies in neonatal hepatocytes, brain microvascular endothelial cells, and NB2a neuronal cells hinted at conservation of this pathway in different cell types [25], [26], other studies have since questioned the role of βOHB as a potential physiological HDAC inhibitor [27], [28]. Interestingly, recent studies in hepatocytes, cortical neurons, myotubes and endothelial cells suggested that βOHB may serve as a novel substrate for a transcriptionally-activating histone modifications. This so called lysine β-hydroxybutyrylation was found in proximity to fasting-relevant hepatic pathways, including amino acid catabolism, circadian rhythm, and PPAR signaling [28], and was found to regulate expression of BDNF [29] and hexokinase 2 [27]. How histones become β-hydroxybutyrylated remains unknown. A series of biochemical experiments suggest that SIRT3 facilitates the de-β-hydroxybutyrylation of histones [30]. While there is thus some evidence to suggest that βOHB may serve as a direct signaling molecule regulating genes, the potency and importance of βOHB as regulator of gene expression in various cell types is unclear. Accordingly, here we aimed to investigate the capacity of βOHB to regulate gene expression and thereby serve as a direct signaling molecule during the fasted state. To this end, we investigated whether βOHB, in comparison to the well-established HDAC inhibitor butyrate, influences in vitro differentiation of adipocytes, macrophages, myotubes. In addition, we studied the effect of βOHB on whole genome gene expression in primary mouse adipocytes, macrophages, myotubes and hepatocytes via RNA-seq.

## MATERIALS AND METHODS

### Materials

βOHB was (R)-(–)-3-Hydroxybutyric acid sodium salt from Sigma-Aldrich (#298360). Butyrate was Sodium butyrate from Sigma-Aldrich (#303410).

### Differentiation experiments

3T3-L1 fibroblasts were maintained in DMEM supplemented with 10% newborn calf serum (NCS) and 1% penicillin/streptomycin (P/S) (all Lonza). Experiments were performed in six-well plates. For Oil red O stainings, cells were differentiated using the standard protocol. Two days post-confluence, cells were switched to DMEM supplemented with 10% fetal bovine serum (FBS), 1% P/S, 0.5 mM isobutylmethylxanthine, 1 µM dexamethasone, 5 µg/ml insulin for 2 days in the presence of either 8 mM βOHB or 8 mM Butyrate. After 2 days cells were switched to DMEM supplemented with 10% fetal bovine serum (FBS), 1% P/S, 5 µg/ml insulin and the tested compounds for another 2 days. Then cells were maintained in normal DMEM medium (2-3 days), in the presence of the tested compounds until the ORO stainings on day 10. For qPCR experiments cells were differentiated using the mild protocol, which allows for more sensitive assessment of compounds promoting the differentiation process at day 4 of differentiation [31]. Two days post-confluence, cells were switched to DMEM supplemented with 10% fetal bovine serum (FBS), 1% P/S, 0.5 mM isobutylmethylxanthine, 0.5 µM dexamethasone, 2 µg/ml insulin for 2 days, with the addition of either 1 μM Rosi, 8 mM βOHB or 8 mM Butyrate. After 2 days, the medium was changed to DMEM supplemented with 10% fetal bovine serum (FBS), 1% P/S, 2 µg/ml insulin and the tested compounds for another 2 days, before cells were harvested for RNA isolation. C2C12 skeletal muscle cells were cultured in DMEM supplemented with 20% FBS (growth medium, GM) and induced to differentiate with DMEM supplemented with 2% horse serum (differentiation medium, DM) upon reaching confluence in the presence of either 5 mM βOHB or 5 mM Butyrate. DM was renewed every other day. Myotube formation was complete (visually) by day 5.

THP-1 cells were cultured in RPMI 1640 + heat-inactivated FBS and 1% P/S. Differentiation to macrophages was induced with 62.5 ng/mL phorbol 12-myristate 13-acetate (PMA, Sigma) for 24 h in the presence of either 8 mM βOHB or butyrate. Microscopic pictures were taken and cells were subsequently frozen for RNA isolation. All cells were cultured at 37 °C with 5% CO_2_.

### Isolation and differentiation of stromal vascular fraction

Inguinal white adipose tissue from 3-4 WT-C57Bl/6 male mice was collected and placed in Dulbecco’s modified eagle’s medium (DMEM; Lonza) supplemented with 1% Penicillin/Streptomycin (PS) and 1% bovine serum albumin (BSA; Sigma-Aldrich). Material was minced finely with scissors and digested in collagenase-containing medium (DMEM with 3.2 mM CaCl_2_, 1.5 mg/ml collagenase type II (C6885, Sigma-Aldrich), 10% FBS, 0.5% BSA, and 15 mM HEPES) for 1 h at 37°C, with occasional vortexing. Cells were filtered through a 100-μm cell strainer (Falcon). Subsequently, the cell suspension was centrifuged at 1600 rpm for 10 min and the pellet was resuspended in erythrocyte lysis buffer (155 mM NH_4_Cl, 12 mM NaHCO_3_, 0.1 mM EDTA). Upon incubation for 2 min at room temperature, cells were centrifuged at 1200 rpm for 5 min and the pelleted cells were resuspended in DMEM containing 10% fetal bovine serum (FBS) and 1% PS (DMEM/FBS/PS) and plated. Upon confluence, the cells were differentiated according to the protocol as described previously [32], [33]. Briefly, confluent SVFs were plated in 1:1 surface ratio, and differentiation was induced 2 days afterwards by switching to a differentiation induction cocktail (DMEM/FBS/PS, 0.5 mM isobutylmethylxanthine, 1 μM dexamethasone, 7 μg/ml insulin and 1 µM rosiglitazone) for 3 days. Subsequently, cells were maintained in DMEM/FBS/PS, and 7 μg/ml insulin for 3-6 days and switched to DMEM/FBS/PS for 3 days. Average rate of differentiation was at least 80% as determined by eye.

### Isolation and differentiation of bone marrow derived monocytes

Bone marrow cells were isolated from femurs of WT-C57Bl/6 male mice following standard protocol and differentiated into macrophages (bone marrow-derived macrophages, BMDMs) in 6-8 days in DMEM/FBS/PS supplemented with 20% L929-conditioned medium (L929). After 6-8 days, non-adherent cells were removed, and adherent cells were washed and plated in 12-well plates in DMEM/FBS/PS + 10% L929. After 24 hours, medium was switched to 2% L929 in DMEM/FBS/PS overnight. Cells were treated the following day.

### Isolation and differentiation of skeletal myocytes

Myoblasts from hindlimb muscle of WT-C57Bl/6 male mice were isolated as previously described [34]. In brief, the muscles were excised, washed in 1× PBS, minced thoroughly, and digested using 1.5 mL collagenase digestion buffer (500 U/ml or 4 mg/mL collagenase type II (C6885, Sigma-Aldrich), 1.5 U/ml or 5 mg/mL Dispase II (D4693, Sigma-Aldrich), and 2.5 mM CaCl2 in 1× PBS) at 37°C water bath for 1 h in a 50 ml tube, agitating the tube every 5 min. After digestion, the cell suspension containing small pieces of muscle tissue was diluted in proliferation medium (PM: Ham’s F-10 Nutrient Mix (#31550023, Thermo Fisher Scientific) supplemented with 20% fetal calf serum, 10% HS, 0.5% sterile filtered chicken embryo extract (#092850145, MP Biomedicals), 2.5 ng/ml basic fibroblast growth factor (#PHG0367, Thermo Fisher Scientific), 1% gentamycin, and 1% PS), and the suspension was seeded onto matrigel-coated (0.9 mg/ml, #354234, Corning) T150 flasks at 20% surface coverage. Cells were grown in 5% CO2 incubator at 37°C. Confluence was reached latest after 5 d in culture, upon which cells were trypsinized (0.25% trypsin), filtered with 70 µm filters, centrifuged at 300 g for 5 min, and then seeded on an uncoated T150 flask for 45 min to get rid of fibroblasts. Subsequently, myoblasts were seeded in PM at 150.000 cells/mL onto matrigel-coated 12-well plates cells. Upon reaching confluence, differentiation was induced by switching to differentiation medium (DM: Ham’s F-10 Nutrient Mix supplemented with 5% horse serum (HS) and 1% PS). DM was replaced every other day. Myotubes fully differentiated by Day 5 of differentiation in DM. The medium was renewed every other day.

### Isolation and culturing of hepatocytes

Primary hepatocytes were isolated from C57BL/6NHsd male mice via collagenase perfusion as described previously [35]–[38]. Cells were plated onto collagen (0.9 mg/ml) coated 24-well plates at 200,000 cells/well in Williams E medium (PAN Biotech), substituted with 10% FBS, 100 nM dexamethasone and penicillin/streptomycin and maintained at 37 °C in an atmosphere with 5% CO2. After four hours of attachment, cells were washed with phosphate buffer saline (PBS) and allowed to rest in dexamethasone-free medium overnight before treatment.

### Treatments

Primary cells were treated for 6h with 5 mM βOHB or Butyrate, with PBS as control. Adipocytes and Macrophages were treated in DMEM/FCS/PS. Myotubes were treated in DM. Hepatocytes were treated in Williams E medium. Cells were washed with PBS once and stored in-80 °C until RNA was isolated.

### RNA isolation & RNA sequencing

Total RNA from all cell culture samples was extracted using TRIzol reagent (Thermo Fisher Scientific) and purified using the Qiagen RNeasy Mini kit (Qiagen) according to manufacturer’s instructions. RNA concentration was measured with a Nanodrop 1000 spectrometer and RNA integrity was determined using an Agilent 2100 Bioanalyzer with RNA 6000 microchips (Agilent Technologies). Library construction and RNA sequencing on BGISEQ-500 were conducted at Beijing Genomics Institute (BGI) for pair-end 150bp runs. At BGI, Genomic DNA was removed with two digestions using Amplification grade DNAse I (Invitrogen). The RNA was sheared and reverse transcribed using random primers to obtain cDNA, which was used for library construction. The library quality was determined by using Bioanalyzer 2100 (Agilent Technologies). Then, the library was used for sequencing with the sequencing platform BGISEQ-500 (BGI). All the generated raw sequencing reads were filtered, by removing reads with adaptors, reads with more than 10% of unknown bases, and low quality reads. Clean reads were then obtained and stored as FASTQ format.

The RNA-seq reads were then used to quantify transcript abundances. To this end the tool *Salmon* [39] (version 1.2.1) was used to map the reads to the GRCm38.p6 mouse genome assembly-based transcriptome sequences as annotated by the GENCODE consortium (release M25) [40]. The obtained transcript abundance estimates and lengths were then imported in R using the package *tximport* (version 1.16.1) [41], scaled by average transcript length and library size, and summarized on the gene-level. Such scaling corrects for bias due to correlation across samples and transcript length, and has been reported to improve the accuracy of differential gene expression analysis [41]. Differential gene expression was determined using the package *limma* (version 3.44.3) [42] utilizing the obtained scaled gene-level counts. Briefly, before statistical analyses, nonspecific filtering of the count table was performed to increase detection power [43], based on the requirement that a gene should have an expression level greater than 20 counts, i.e. 1 count per million reads (cpm) mapped, for at least 6 libraries across all 36 samples. Differences in library size were adjusted by the trimmed mean of M-values normalization method [44]. Counts were then transformed to log-cpm values and associated precision weights, and entered into the *limma* analysis pipeline [45]. Differentially expressed genes were identified by using generalized linear models that incorporate empirical Bayes methods to shrink the standard errors towards a common value, thereby improving testing power [42], [46]. Genes were defined as significantly changed when P < 0.001.

### Biological interpretation of transcriptome data RNA isolation & RNA sequencing

Changes in gene expression were related to biologically meaningful changes using gene set enrichment analysis (GSEA) [47]. It is well accepted that GSEA has multiple advantages over analyses performed on the level of individual genes [47]–[49]. GSEA evaluates gene expression on the level of gene sets that are based on prior biological knowledge, e.g. published information about biochemical pathways or signal transduction routes, allowing more reproducible and interpretable analysis of gene expression data. As no gene selection step (fold change and/or p□value cut-off) is used, GSEA is an unbiased approach. A GSEA score is computed based on all genes in gene set, which boosts the signal-to-noise ratio and allows to detect affected biological processes that are due to only subtle changes in expression of individual genes. This GSEA score called normalized enrichment score (NES) can be considered as a proxy for gene set activity. Gene sets were retrieved from the expert-curated KEGG pathway database [50]. Only gene sets comprising more than 15 and fewer than 500 genes were taken into account. For each comparison, genes were ranked on their t□value that was calculated by the moderated t□test. Statistical significance of GSEA results was determined using 10,000 permutations.

### Statistical analyses

Statistical analysis of the transcriptomics data was performed as described in the previous paragraph. Data are presented as mean ± SD. P-values < 0.05 were considered statistically significant.

## RESULTS

### Butyrate but not β-hydroxybutyrate impairs differentiation of adipocyte, monocyte and macrophage cell lines

To solidify the concept of βOHB being a powerful signaling molecule that influences cellular homeostasis, we examined whether βOHB affects cellular differentiation. Previously, we showed that butyrate, despite acting as a selective PPARγ agonist, inhibits adipogenesis in 3T3-L1 cells [51]. Due to structural and possibly functional resemblance with butyrate, we hypothesized that βOHB might exert similar effects on the differentiation of 3T3-L1 cells. Compared to the control, 8 mM βOHB did not visibly affect adipocyte differentiation, as assessed during the differentiation process (Day 4) and terminally (Day 10; Figure 1A). By contrast and in line with previous studies, 1 µM rosiglitazone stimulated the differentiation process (Day 4), whereas 8 mM butyrate markedly inhibited adipocyte differentiation (Day 4 and 10; Figure 1A). Corroborating the visual assessment, rosiglitazone significantly induced expression of the adipogenic marker genes *Adipoq, Slc2a4 (Glut4)* and *Fabp4*, whereas butyrate significantly downregulated these genes. In line with the lack of effect on 3T3-L1 differentiation, βOHB had a minor impact on the expression of *Slc2a4 (Glut4)* and no impact on the expression of *Adipoq* or *Fabp4* (Figure 1B).

**Figure 1:**
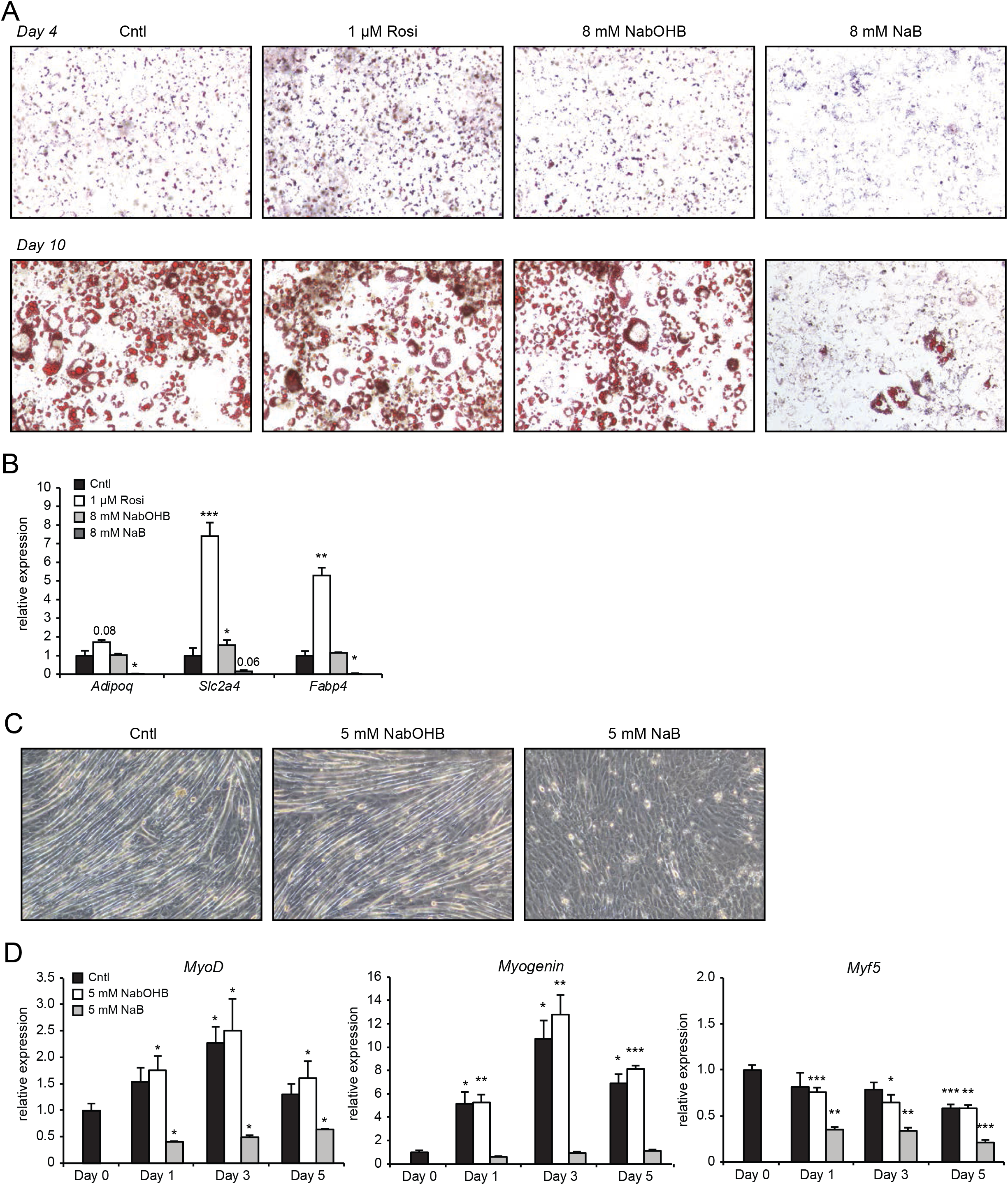
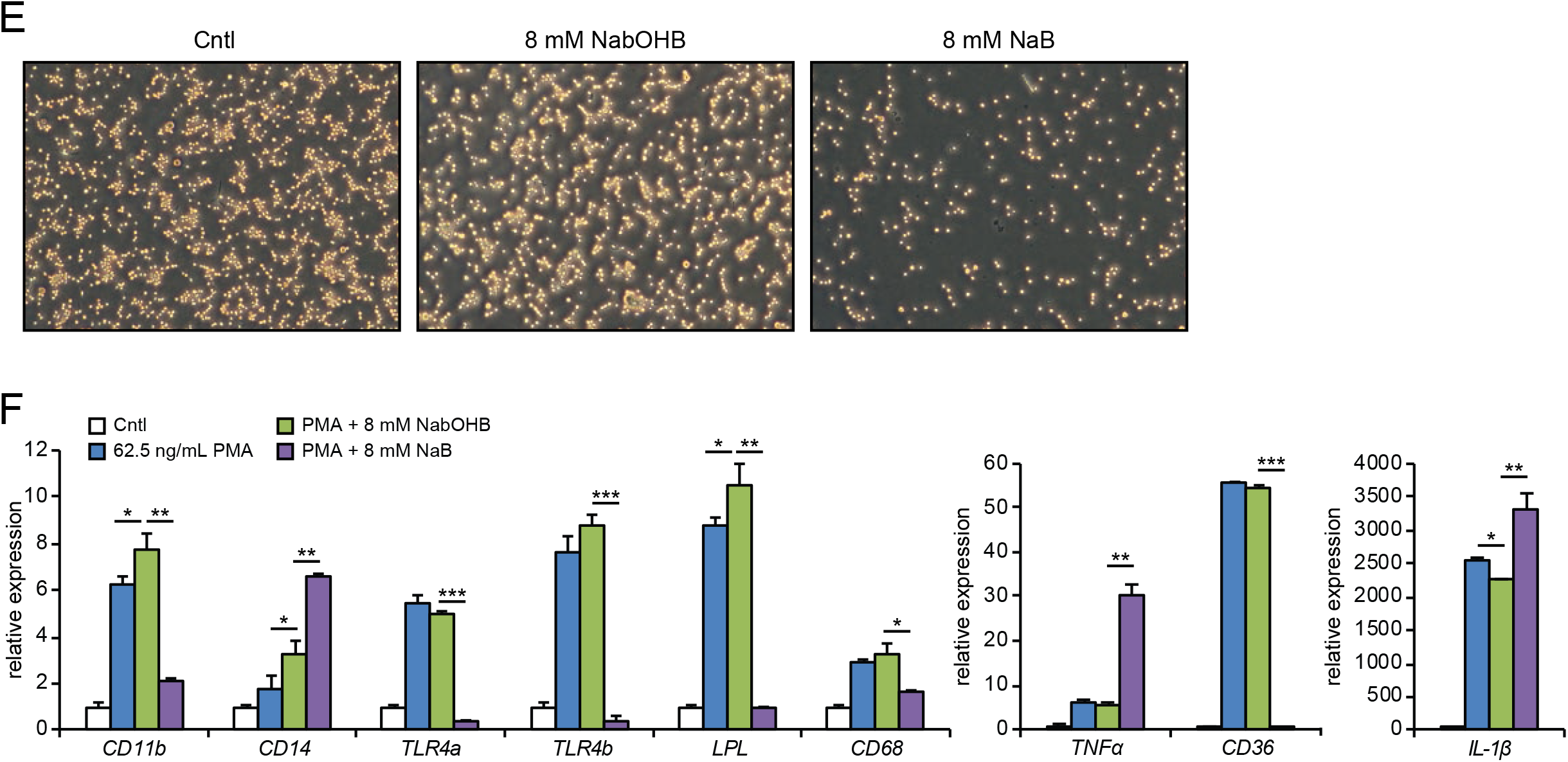
Differential effects of βOHB and butyrate on the differentiation process of 3T3L1 adipocytes and C2C12 myotubes. (A) Representative Oil red O staining of 3T3-L1 adipocytes at day 4 of the standard differentiation protocol. (B) Expression of differentiation markers and PPARγ targets determined by qPCR at day 4 using the mild differentiation protocol in the presence of either 1 μM Rosi, 8 mM βOHB or 8 mM butyrate. (C) Representative microscopic pictures of myotube formation after 5 days of differentiation in the presence of 5 mM βOHB or 5 mM butyrate. (D) Gene expression of myocyte differentiation markers *MyoD, Myogenin* and *Myf5* after differentiation. (E) Representative pictures of THP-1 cells differentiating for 24h in 62.5 ng/mL PMA in presence of either 8 mM βOHB or 8 mM butyrate. (F) Gene expression of macrophage differentiation markers. Asterisks indicate significant differences according to Student’s t test (* p < 0.05; ** p < 0.01; *** p < 0.001).

Next we studied myogenesis. Butyrate was previously reported to inhibit myogenesis when present during the differentiation process [52]. To assess whether βOHB might influence myogenesis, we differentiated C2C12 myoblasts in the presence of 5 mM βOHB or 5 mM butyrate. In line with previous reports, butyrate inhibited the differentiation of myoblasts towards myotubes (Figure 1C) [52]. By contrast, βOHB did not visibly impact myotube formation (Figure 1C). Myogenesis is driven by muscle regulatory factors (MRFs) including *MyoG, MyoD* and *Myf5* [53], [54]. Supporting the lack of effect of βOHB on myogenesis, expression levels of all three MRFs were similar in βOHB and control-treated C2C12 cells at any time-point during the differentiation process (Figure 1D). This is in clear contrast to treatment with butyrate, which prevented upregulation of *MyoG* and *MyoD* and downregulated *Myf5* at all time-points, respectively. We also wondered whether instead of influencing the differentiation process, βOHB might affect the polarization of myotubes to either myosin heavy chain class I (MHCI) or class II (MHCII). Expression of *Myh3, Myh7* and *Myh8*, representing MHCI, were unchanged between βOHB and control treated myoblasts. Expression of *Myh1, Myh2* and *Myh4*, representing MHCII, were also unchanged between βOHB and control (Supplemental figure 1A), suggesting that βOHB does not influence the polarization of myotubes when added during the differentiation process. Lastly, βOHB as well as butyrate have been reported to modulate immune cell function and viability [22], [55]. Specifically, butyrate demonstrated pro-apoptotic effects on THP-1 in previous studies [56]–[58]. To assess whether either compound influences the differentiation of a monocytic cell line *in vitro*, we differentiated THP-1 cells with PMA in the presence of 8 mM βOHB or butyrate. Corroborating reports of pro-apoptotic effects of butyrate on THP-1 cells [56]–[58], addition of butyrate during the differentiation process resulted in a clear reduction in the density of monocytes (Figure 1E). In keeping with the lack of effect on myocyte and adipocyte differentiation, βOHB also did not visually impact on THP-1 cell differentiation (Figure 1E). PMA-induced differentiation of THP-1 cells is marked by differential expression of several marker genes including *CD11b, CD14, TNF-*α and *CD68* [59]–[62]. Butyrate prevented PMA-mediated induction of *CD11b* and *CD68*, while further inducing *TNF-*α, *CD14*, and *IL-1*β expression (Figure 1F). In addition, butyrate markedly suppressed the expression of the pattern recognition receptor *TLR4a* and *TLR4b* and the lipid-associated genes *LPL* and *CD36*. By contrast, gene expression changes by βOHB for most genes were non-significant relative to cells treated with PMA only (Figure 1F). Interestingly, βOHB significantly altered gene expression of *CD11b, CD14, LPL* and *IL-1*β, although the magnitude of the effect was very modest (Figure 1F). These results suggest that butyrate exerts a strong effect on the differentiation and viability of THP-1 cells. In comparison, the effects of βOHB are small.

### β-hydroxybutyrate alters gene expression in primary myocytes but not primary adipocytes, macrophages, and hepatocytes

We reasoned that if βOHB has a signaling function, it would likely alter the expression of genes either directly or indirectly. Accordingly, we investigated the ability of βOHB to regulate gene expression in cells that have been suggested to be targeted by βOHB. Specifically, we collected primary mouse adipocytes, primary mouse bone-marrow derived macrophages, primary mouse myotubes, and primary mouse hepatocytes and performed RNA-sequencing after 6h treatment with either 5 mM βOHB and 5 mM butyrate. Importantly, the RNAseq data showed that all cell types expressed at least one type of the monocarboxylate transporters *Slc16a1* (*Mct1), Slc16a7* (*Mct2)* and *Slc16a6* (*Mct7)*, that are responsible for the transport of βOHB and butyrate [22], [63]–[65] (Figure 2A).

**Figure 2:**
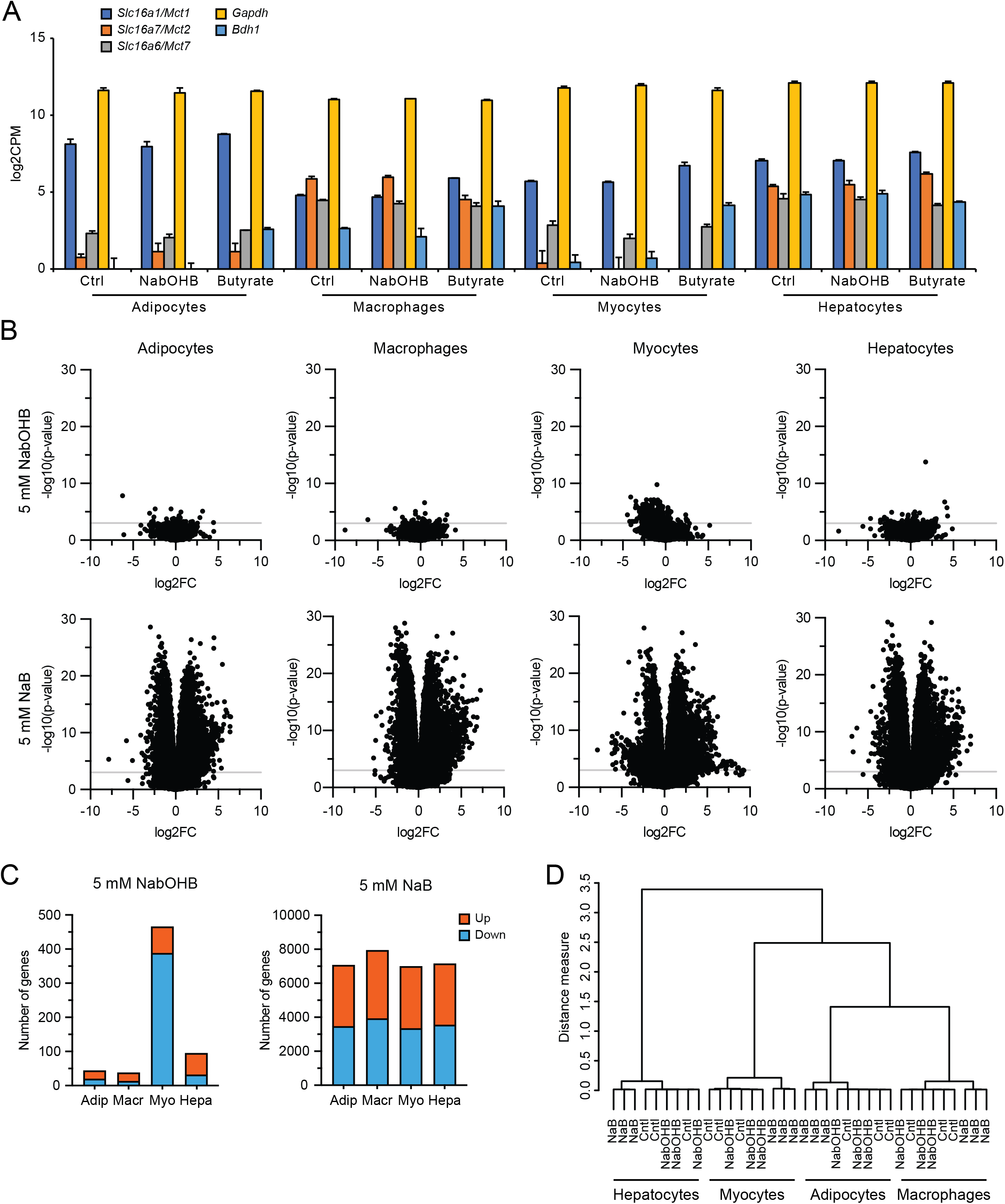
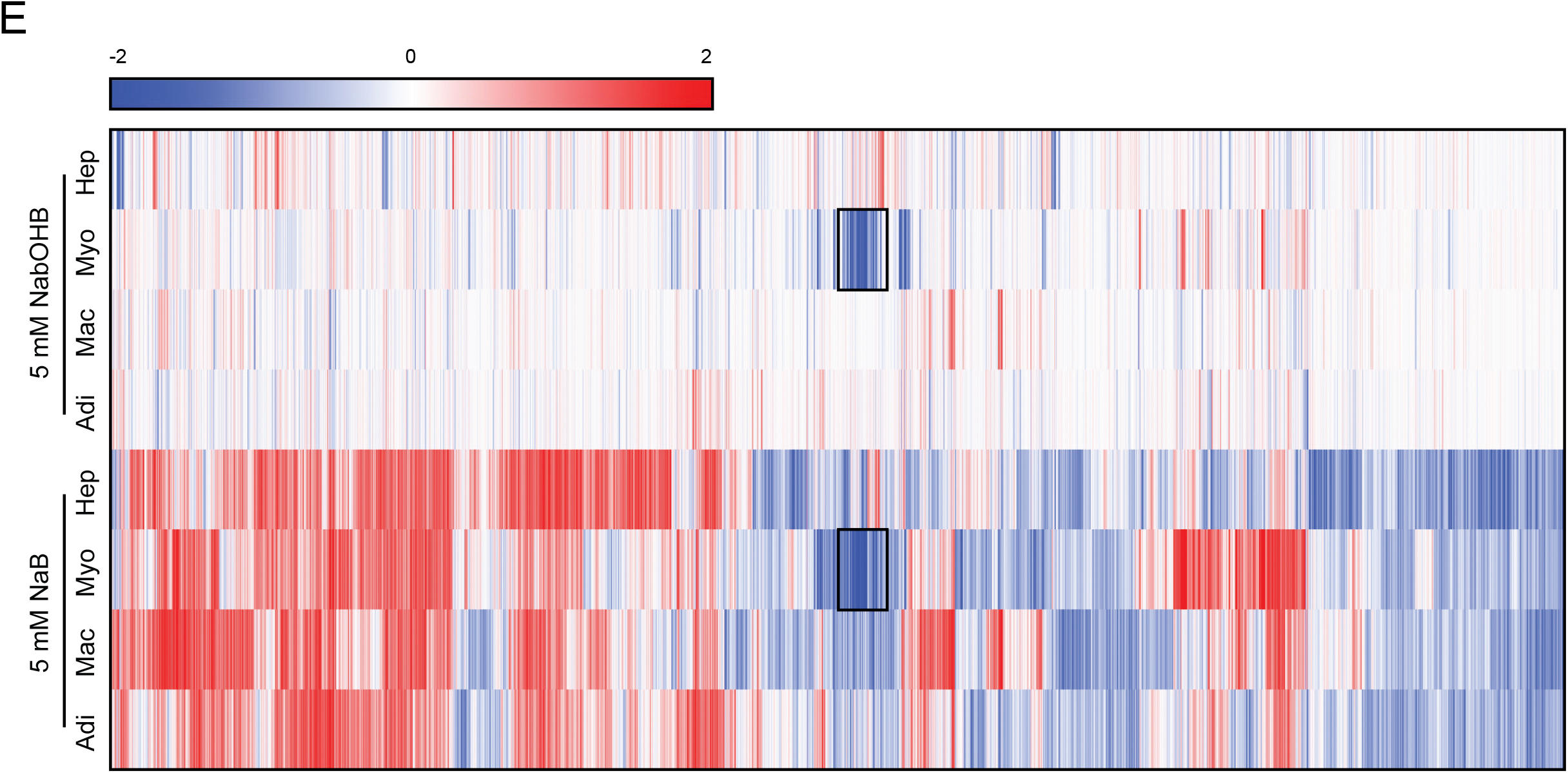
Disparate effects of βOHB and butyrate on gene expression in primary adipocytes, macrophages, myocytes and hepatocytes. (A) Expression levels (log2CPM) of monocarboxylate transporters *Mct1, Mct2* and *Mct7* in relation to *Gapdh* and *Bdh1*. (B) Volcano plots showing log2[fold-change] (x-axis) and the-10log of the raw p-value (y-axis) for every cell type treated with βOHB and butyrate. The grey line indicates p = 0.001. (C) Number of genes significantly (p < 0.001) altered by treatment with βOHB and butyrate. (D) Hierarchical clustering of βOHB and butyrate treated samples. (E) Hierarchical biclustering of βOHB and butyrate treated samples visualized in a heatmap. Clustered are significant differentially expressed genes based on Pearson correlation with average linkage. Red indicates upregulated, blue indicates downregulated. Black reactangle marks genes that appear similarly regulated by βOHB and butyrate in myocytes.

The cells treated with butyrate showed an anti-conservative p-value distribution, suggesting that butyrate has a marked effect on gene expression in all cell types studied. Conversely, cells treated with βOHB showed a uniform or conservative p-value distribution (Supplemental figure 2A), suggesting that βOHB treatment minimally impacted gene expression. To study the magnitude of gene regulation by βOHB and butyrate in the various primary cells, we performed Volcano plot analysis. Strikingly, the effect of βOHB on gene expression was very limited in all cell types, with only a small number of genes reaching the statistical threshold of p < 0.001 (Figure 2B). Using this statistical threshold, βOHB significantly altered expression of 44, 38, 466 and 95 genes in adipocytes, macrophages, myocytes and hepatocytes, respectively. Of these genes, 20, 13, 388 and 32 were downregulated, respectively (Figure 2C). In adipocytes, macrophages, and hepatocytes, less than 10 genes had a false discovery q-value below 0.05, indicating that most of the significant genes in these cells likely represent false positives. In myocytes, 560 genes had a FDR q-value below 0.05 (Supplemental figure 2B). In stark contrast to the relatively minor effects of βOHB, butyrate had a huge effect on gene expression in all primary cells (Figure 2B). Butyrate significantly changed the expression of 7068, 7943, 6996 and 7158 genes in adipocytes, macrophages, myocytes and hepatocytes, respectively (p < 0.001), of which 50 – 52% were downregulated (Figure 2C). The number of differentially expressed genes is similarly high when using a FDR q-value of 0.05 (Supplemental figure 2B).

To further examine the overall effect of βOHB and butyrate on gene regulation in the various cell types, we performed hierarchical clustering and principle component analysis. Both analyses showed that the samples cluster by cell type first. Whereas the butyrate-treated samples clustered apart from the control and βOHB samples in each cell type, the control and βOHB samples did not cluster separately from each other (Figure 2D,E, Supplemental figure 2C). Collectively, these data indicate that in comparison to butyrate, βOHB minimally impacted gene expression in adipocytes, macrophages, and hepatocytes. By contrast, βOHB had a more pronounced effect on gene expression in myocytes, although still much less than observed for butyrate.

### Significant overlap in gene regulation by butyrate across various cell types

Next, we studied the similarity in gene regulation by butyrate among the different cell types. Hierarchical biclustering of all significantly regulated genes per condition showed marked similarity in the response to butyrate. Furthermore, Venn diagrams for the butyrate-treated cells revealed that a large fraction of the significantly regulated genes were shared in all cell types, confirming the similarity in gene regulation by butyrate. Indeed, 18% (1250 genes) of all significantly upregulated genes were upregulated in every cell type. Similarly, 15% (1095 genes) of all significantly downregulated genes were downregulated in every cell type (Figure 3A). Heatmaps of the TOP20 most significantly regulated genes by butyrate showed comparable signal log ratios in all 4 cell types (Figure 3B). Interestingly, the heatmaps for butyrate lists several genes related to histone metabolism (*H1f0, H1f2, H1f4, H1f3, Hcfc1, Phf2, Anp32b*).

**Figure 3:**
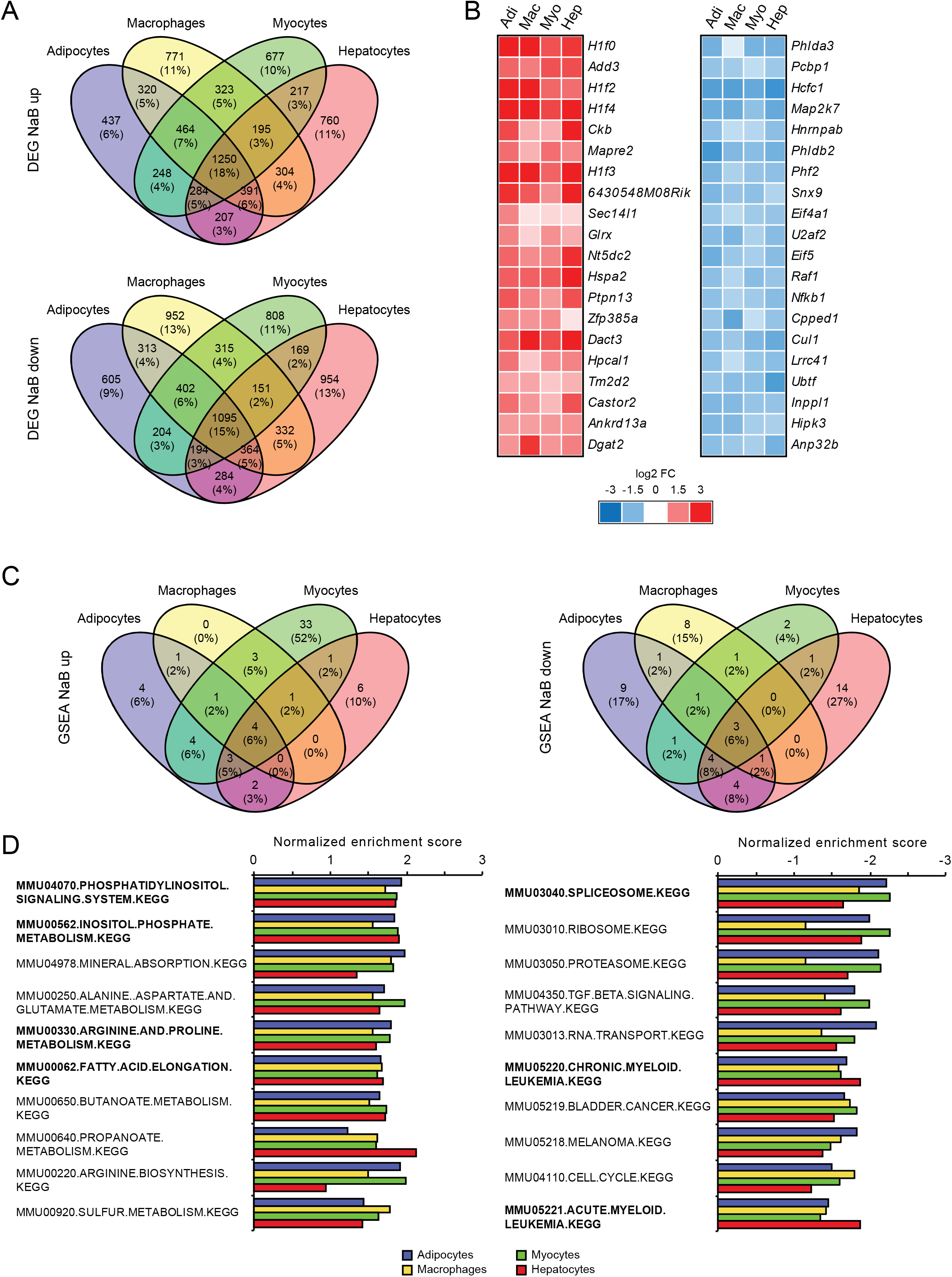
Consistency of gene expression changes elicited by butyrate. (A) Venn diagrams showing overlap in significantly regulated genes by butyrate between cell types (p < 0.001), separated into up- and downregulated genes (B) Heatmaps showing genes that are significantly regulated in all 4 cell types (butyrate). Genes with thick black border are significantly regulated (p < 0.001). In butyrate-heatmaps all listed genes are significantly regulated and genes are sorted by significance. Top 10 up- and downregulated gene sets in βOHB (C) and butyrate-treated cells (D). Gene sets were determined by gene set enrichment analysis (GSEA) based on t-values for the top 100 up- and downregulated genes and are ranked according to averaged normalized enrichment score (NES). Pathways in bold are significantly enriched in all 4 cell types.

To examine the similarity in gene regulation by butyrate across the different cell types at the level of pathways, we performed GSEA analysis using the TOP100 up and down regulated genes according to the T-statistic. The overlap in significantly regulated pathways (FDR q < 0.1) are shown in a Venn diagram, revealing a high overlap for butyrate-induced and repressed pathways among the 4 cell types. 20 out of 61 pathways were induced in at least 2 cell types, while 4 pathways (“phosphatidylinositol-signaling-system”, “62-inositol-phosphate-metabolism”, “arginine-and-proline-metabolism”, and “-fatty-acid-elongation”) were induced in all 4 cell types (Figure 3C). Interestingly, 33 pathways were exclusively induced by butyrate in myocytes (Figure 3C). Conversely, 19 out of 52 pathways were repressed in at least 2 cell types, while the 3 pathways “spliceosome”, “chronic-myeloid-leukemia” and “bladder-cancer”) were repressed in all 4 cell types (Figure 3C). Plotting the TOP10 induced and repressed pathways by average NES scores corroborates the consistent regulation of pathways by butyrate among the various cell types (Figure 3D). Collectively, these analyses indicate considerable overlap in the effect of butyrate on gene expression in all cell types at the gene and pathway level.

### Significant effect of βOHB on gene regulation in primary myocytes

Given the minimal number of genes altered by βOHB in adipocytes, macrophages and hepatocytes, most likely representing false positives, we did not further perform any analyses for these cell types. Instead, we focused our attention on the effects of βOHB on gene regulation in myocytes. Having noted a region of overlap between βOHB and butyrate (Figure 2E; black rectangle), we first investigated the similarity in gene regulation between both compounds in myocytes. Venn diagram analysis revealed that of the 451 genes downregulated by βOHB according to FDR q<0.05, 320 genes (71%) were also significantly downregulated by butyrate. Likewise, 50% of the 109 genes upregulated by βOHB were also upregulated by butyrate (Figure 4A). Supplemental table 1 shows a list of genes regulated by βOHB according to FDR p<0.001.

**Figure 4:**
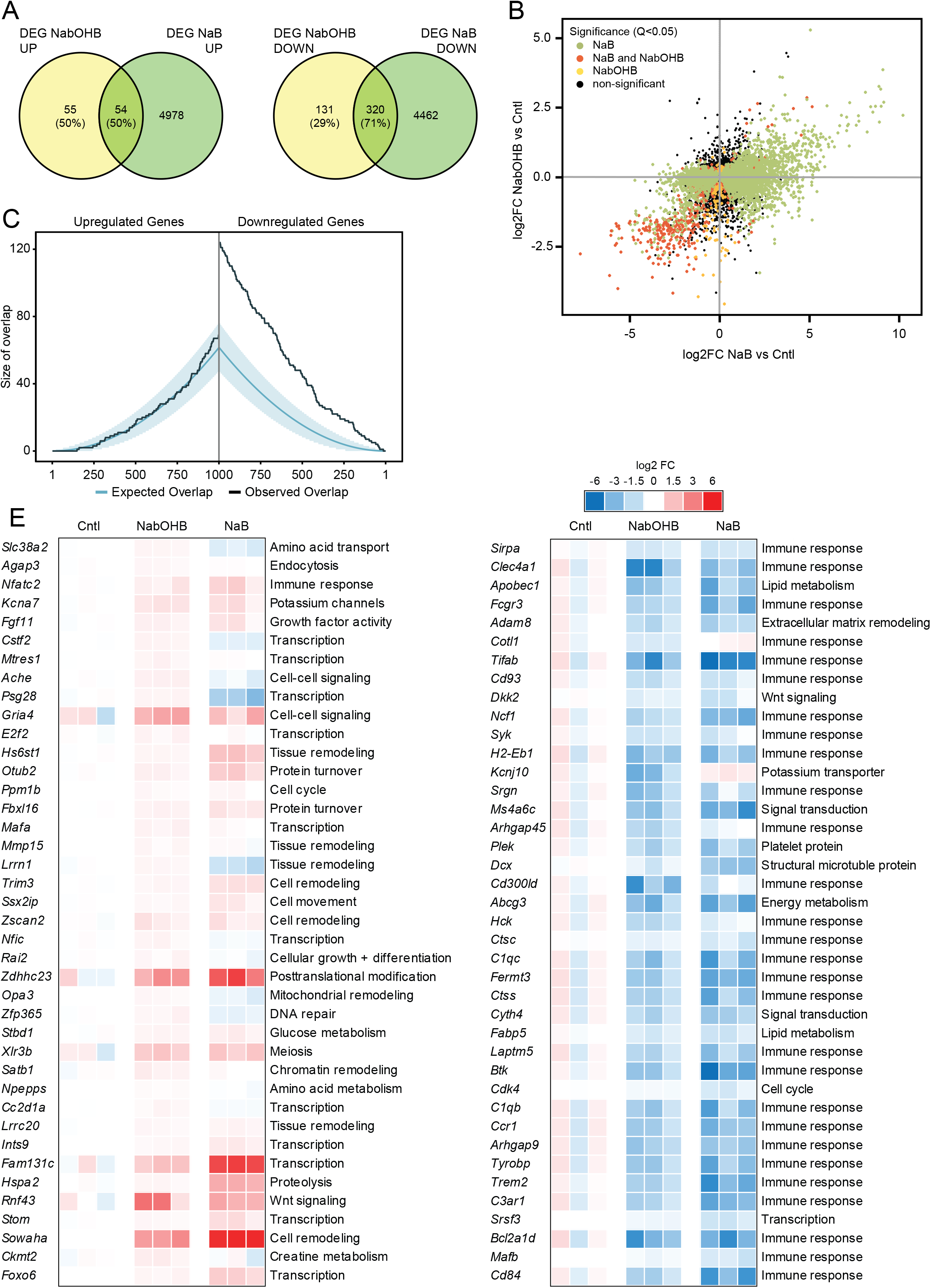
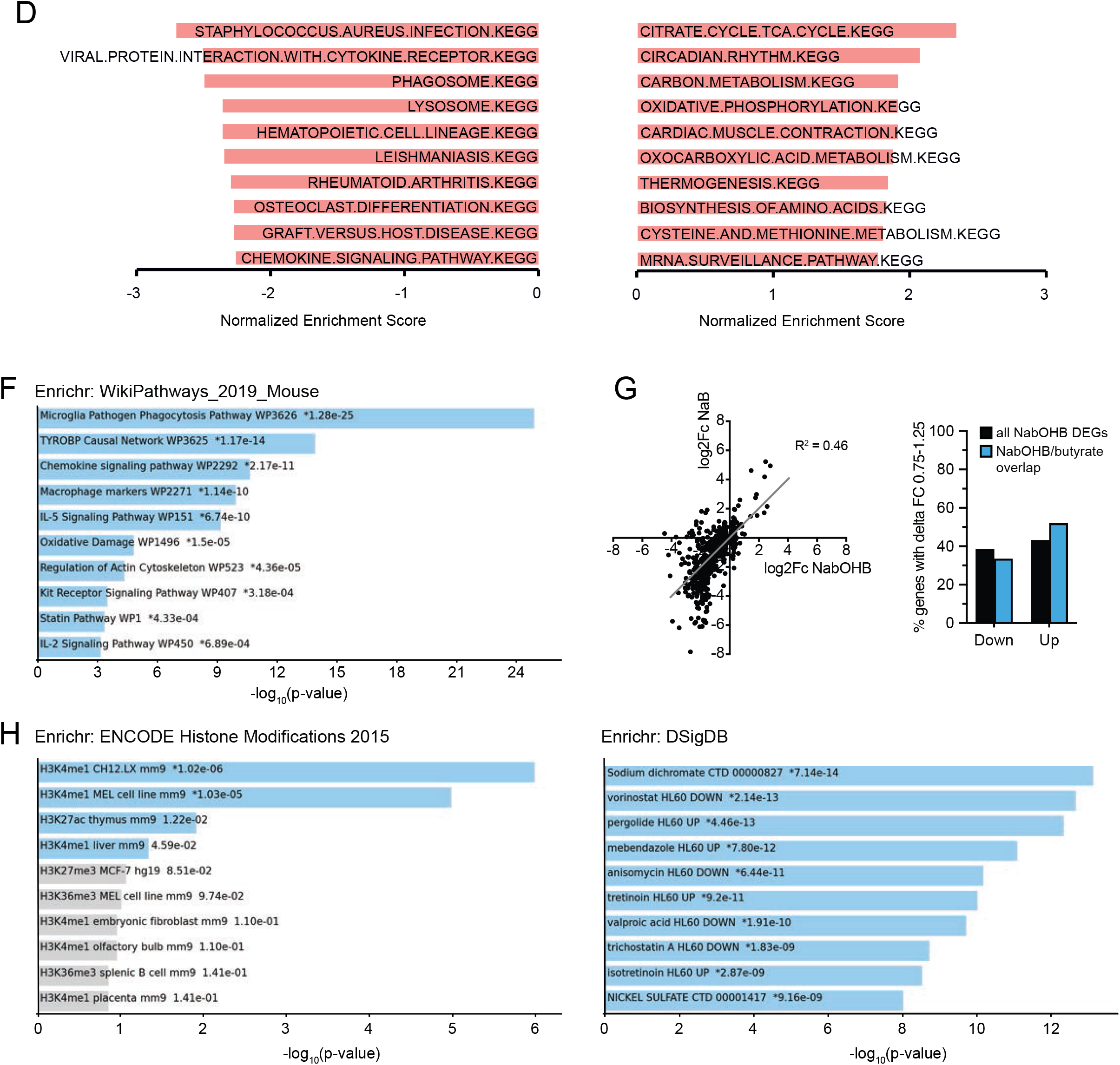
βOHB regulates genes and pathways related to TCA cycle and Immunity in primary myocytes. (A) Venn diagrams showing overlap of significantly regulated genes between of βOHB and butyrate regulates genes in myocytes (p < 0.001). (B) Correlation plot of the genes regulated by βOHB. (C) Overlap plot depicting the size of the overlap for genes upregulated (left) or downregulated (right) by βOHB and butyrate treatment. The size of the overlap for randomly selected gene sets is shown by the blue line (blue shading depicts confidence interval). The observed overlap is shown by the black line. (D) Gene sets negatively enriched for βOHB treatment in myocytes according to GSEA using TOP100 up- and downregulated genes (FDR q-value < 0.1). Gene sets are ranked according to Normalized Enrichment Score. (E) Heatmaps showing TOP40 up- and downregulated genes by βOHB in primary myotubes, alongside butyrate. (F) Enrichr analysis of βOHB downregulated genes (p<0.001) according to ‘WikiPathways 2019 mouse’. (G) Quantitation of genes that βOHB or βOHB and butyrate (overlap) regulated more similarly (FC ratio between 0.75x and 1.25x) split into up and down-regulated genes. (H) Enrichr analysis of βOHB regulated genes (p<0.001) according to ‘ENCODE Histone modifications’ and ‘DSigDB’. Significance in Enrichr analyses: Blue bars indicate p < 0.05. An asterisk (*) next to a p-value indicates the term also has a significant adjusted p-value (<0.05).

To further examine the similarity in gene regulation between βOHB and butyrate, we plotted log2Fc values of all genes in a correlation plot. The correlation plot showed considerable overlap in gene regulation between βOHB and butyrate, which was most obvious for the genes downregulated by the two treatments (Figure 4B). To statistically analyze the overlapping gene regulation, we performed overlap analysis [66], [67]. In this analysis, the expected overlap is calculated for any number of top genes (on the x-axis) using a hypergeometric distribution (i.e. over-representation analysis). The blue line and shaded blue area cover the expected overlap under the null hypothesis (95% CI), the black line indicates the observed overlap (Figure 4C). Consistent with the Venn diagram and scatter plot, significant overlap was observed between βOHB and butyrate for the downregulated genes but not for the upregulated genes. This may indicate a similar mode of action for both compounds.

To gain further insight into the pathways regulated by βOHB in myocytes, we performed gene set enrichment analysis (GSEA) and Enrichr analysis, first focusing on the upregulated pathways. Using a statistical threshold of q<0.1, GSEA yielded 25 gene set that were significantly upregulated by βOHB in myocytes (Figure 4D; Supplementary table 2). Many of the upregulated gene sets were related to metabolic pathways, including the TCA cycle, oxidative phosphorylation, and amino acid metabolism. Enrichr analysis (‘WikiPathways Mouse’) on the 78 upregulated genes that met the statistical significance threshold of p<0.001 yielded only one significant pathway (adjusted p<0.05), which was TCA cycle (not shown). The TOP40 list of most highly upregulated genes presents a diverse set of genes involved in Cell cycle progression, tissue and cell remodeling as well as gene regulation (Figure 4E).

With respect to down regulation of gene expression, using a statistical threshold of q<0.1 for the GSEA analysis, 96 gene sets were significantly downregulated by βOHB in myocytes (Supplementary table 3). Especially the downregulated pathways were related to immunity and inflammation (Figure 4D) and Enrichr analysis (‘WikiPathways Mouse’) confirmed the enrichment in inflammation-related pathways (Figure 4F). The downregulation of genes involved in immunity and inflammation was reflected in the top 40 list of most highly downregulated genes (Figure 4E). The majority of these genes was similarly downregulated by βOHB and butyrate, suggesting a common mechanism of regulation.

Lastly, to substantiate the notion that βOHB and butyrate might affect gene expression via a common mechanism, we plotted log2Fc values for genes significantly downregulated by βOHB in a correlation plot and determined the number of genes that fell within a fold-change ratio of 0.75x to 1.25x. Approximately 40-50% of all βOHB DEGs and genes regulated by βOHB and butyrate fell within this artificial cut-off, indicating that a substantial number of genes regulated by βOHB were regulated by butyrate to a similar extent (Figure 4G). Enrichr analysis of βOHB-downregulated genes for ‘Encode Histone modifications’ and ‘DSigDB’ showed significant overlap with gene signatures belonging to histone modification experiments and treatments with common HDAC inhibitors, including Vorinostat, Valproic acid and Trichostatin A (Figure 4H). These data suggest that, in accordance with butyrate’s well established HDAC inhibitory function [68], βOHB may also regulate target genes via epigenetic mechanisms in primary myocytes.

## 2. DISCUSSION

In this paper we studied the potential of β-hydroxybutyrate (βOHB) to influence cellular differentiation and for the first time performed whole genome expression analysis in primary adipocytes, macrophages, myocytes and hepatocytes comparing βOHB side-by-side with the well-established HDAC inhibitor butyrate. At physiologically relevant plasma concentrations of βOHB as measured after fasting or ketogenic diet, βOHB did not affect the differentiation of 3T3-L1, C2C12 or THP-1 cells. Furthermore, βOHB minimally influenced gene expression in primary adipocytes, macrophages and hepatocytes but altered the expression of a substantial number of genes in primary myocytes. The results from βOHB are in stark contrast to the consistent butyrate-mediated inhibition of differentiation in 3T3-L1, C2C12 and THP-1 cells, and the profound and consistent gene expression changes caused by butyrate in the various primary cells. Together, these data do not support the notion that βOHB serves as a potent signaling molecule regulating gene expression in adipocytes, macrophages and hepatocytes. The suppressive effect of βOHB in myocytes on the expression of genes involved in immunity merits further study.

Interest in ketones has surged in the recent years. Illustrated by the sheer abundance of reviews and perspective papers on the potential benefits of ketosis, βOHB is considered as a potential mediator of putative fasting-related health benefits [2], [12]–[23]. Common to all reviews is the prominent portrayal of βOHB as a potent HDAC inhibitor influencing gene expression, a notion originating from work by Shimazu *et al*. in kidney and HEK293 cells [24]. In this study, evidence was presented that βOHB is an endogenous and specific inhibitor of class I histone deacetylases (HDACs) *in vitro* and *in vivo*, leading to protection against oxidative stress. However, several studies published since then have been unable to confirm a HDAC inhibitory activity for βOHB in various cell types, using butyrate as positive control [27]–[29]. Irrespective of the precise mechanism, epigenetic alterations ultimately require changes in gene expression to impact homeostasis. In our differentiation experiments, co-incubation with βOHB did not alter expression of key differentiation genes in 3T3-L1, C2C12 and THP-1 cells. Further unbiased assessment of whole genome expression in mouse primary cells revealed minimal effects of βOHB on gene expression in adipocytes, macrophages, and hepatocytes. In fact, we suspect that all genes significantly altered by βOHB in these cells represent false positives. Assuming that βOHB is taken up by hepatocytes, adipocytes and macrophages, these results contradict the notion that βOHB acts as a general HDAC inhibitor.

An interesting finding of this study was that βOHB had distinct effects on gene expression in primary myotubes. Supporting the use of βOHB in muscle tissue as a substrate for ATP synthesis [69], [70], pathways related to TCA cycle and oxidative phosphorylation were upregulated by βOHB. Additionally, βOHB markedly influenced immunity-related pathways and specifically downregulated various genes belonging to cytokine and chemokine signal transduction, including *Sirpa, Clec4a1, Fcgr3, Cd93, Syk, Ms4a6c, Hck, C1qc, Btk, C1qb* and *Ccr1*. Considering that the *Mct* transporter expression profile is similar among the primary cells, it is unclear why βOHB only exerted these effects in myocytes and not for example in macrophages. Nevertheless, one could speculate that the downregulation of immune-related pathways in muscle cells by βOHB may be part of a broader mechanism to suppress immunity during starvation. Indeed, it is well recognized that starvation present a trade-off between, on the one hand, saving energy to prolong survival and, on the other hand, investing a sufficient amount of energy to maintain immune defenses. It can be hypothesized that βOHB may serve as a signaling molecule that mediates the suppressive effect of starvation on specific immune-related processes [71], [72]. Interestingly, while not supported by the results in adipocytes, macrophages and hepatocytes, Enrichr analysis does suggest an epigenetic mode of action for βOHB in myocytes. Further studies will need to expand on the tissue-specific effects of βOHB and probe the functional significance of above-mentioned findings with *in vivo* knockout studies.

In contrast to βOHB, the effects of butyrate on gene expression were prominent and displayed consistency between the tested primary cell lines and the differentiation experiments. A significant portion of histone metabolism-related genes were consistently regulated between the various cell types. Additionally, the most highly enriched pathways were significantly enriched in most if not all cells. In line with butyrate’s well-established effects on gene expression, pathways relevant to transcriptional activities were enriched as well. Additional analyses using Enrichr are in support of butyrate’s prominent HDAC inhibitory action. The marked effect of butyrate effect on adipocyte and myocyte differentiation in 3T3-L1 and C2C12 cells is in line with previous research [51], [52] and may also partly be explained by epigenetic mechanisms [73]. It should be noted, though, that the data presented here are not suitable to deduce potential physiological effects of butyrate and SCFA fermentation *in vivo*. Juxtaposing the supraphysiological concentration of 5 mM employed in this study are reports of 1 – 12 µM and 14 – 64 µM butyrate in the peripheral and central blood circulation measured in sudden death victims [74].

The main limitation of our study is the exclusive utilization of *in vitro* systems. We opted for this approach to allow for the identification of target genes that may be consistently regulated in more than one cell type in a controlled environment. While novel target genes would have to be replicated *in vivo*, this approach seemed more reasonable for this purpose compared to *in vivo* systems in which it is impossible to study the transcriptional regulation specifically attributable to βOHB. For example, the hepatic response to fasting is shaped by the FFA-PPARα axis, which regulates nearly every branch in lipid metabolism and is indispensable for the physiological adaptation to fasting [75], [76]. The increase of ketone body levels during fasting occurs concurrent with many other metabolic and hormonal changes, including increased plasma fatty acids, cortisol, and glucagon levels, and decreased plasma insulin and leptin levels.

In conclusion, this work is the first to systematically assess the potential of the ketone body βOHB to influence gene expression in various primary cell types by RNA-sequencing. With the exception of myocytes, the lack of genes commonly regulated among cell types coupled to generally insignificant effects on gene expression contradict the notion that βOHB serves as a powerful and general signaling molecule regulating gene expression during the fasted state *in vivo*. Instead our data support the idea that βOHB acts as a niche signaling molecule regulating specific pathways in specific tissues, for example muscle. Mechanistically, this action may include gene expression changes potentially via epigenetic effects but could also be secondary to oxidation or receptor activation. Collectively, in our view, the data presented here do not support the current portrayal of βOHB in literature as the do-it-all-substrate during the fasted state and suggest that βOHB’s effects may be much more nuanced and context-specific. Future research is necessary to delineate the role of βOHB including the regulation of gene expression in a tissue/context-specific manner, as for example in muscle tissue.

## Abbreviations

βOHB: β-hydroxybutyrate
AcAc: acetoacetate
Kbhb: lysine β-hydroxybutyrylation
HDAC: histone deacetylase

