## Supplemental figure 1 for "Transcriptomic analysis reveals niche gene expression effects of beta-hydroxybutyrate in primary myotubes"

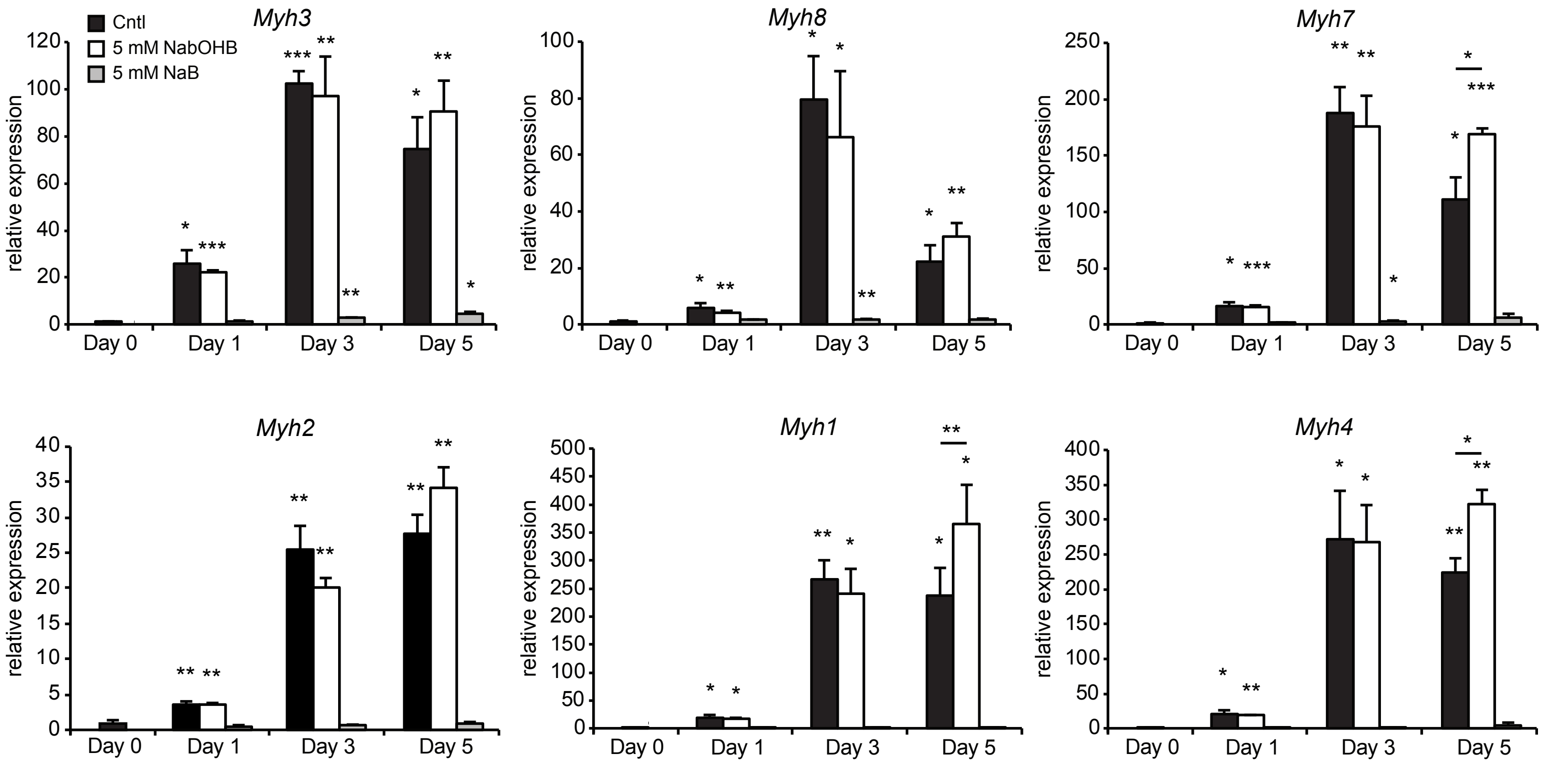


Supplemental figure 1: Gene expression of myotube polarization markers for type I and type II muscle fibers. Error bars represent SD. Asterisks indicate significant differences according to Student's t test (* $p < 0.05$; ** $p < 0.01$; *** $p < 0.001$).
