## Supplemental figure 2 for "Transcriptomic analysis reveals niche gene expression effects of beta-hydroxybutyrate in primary myotubes"

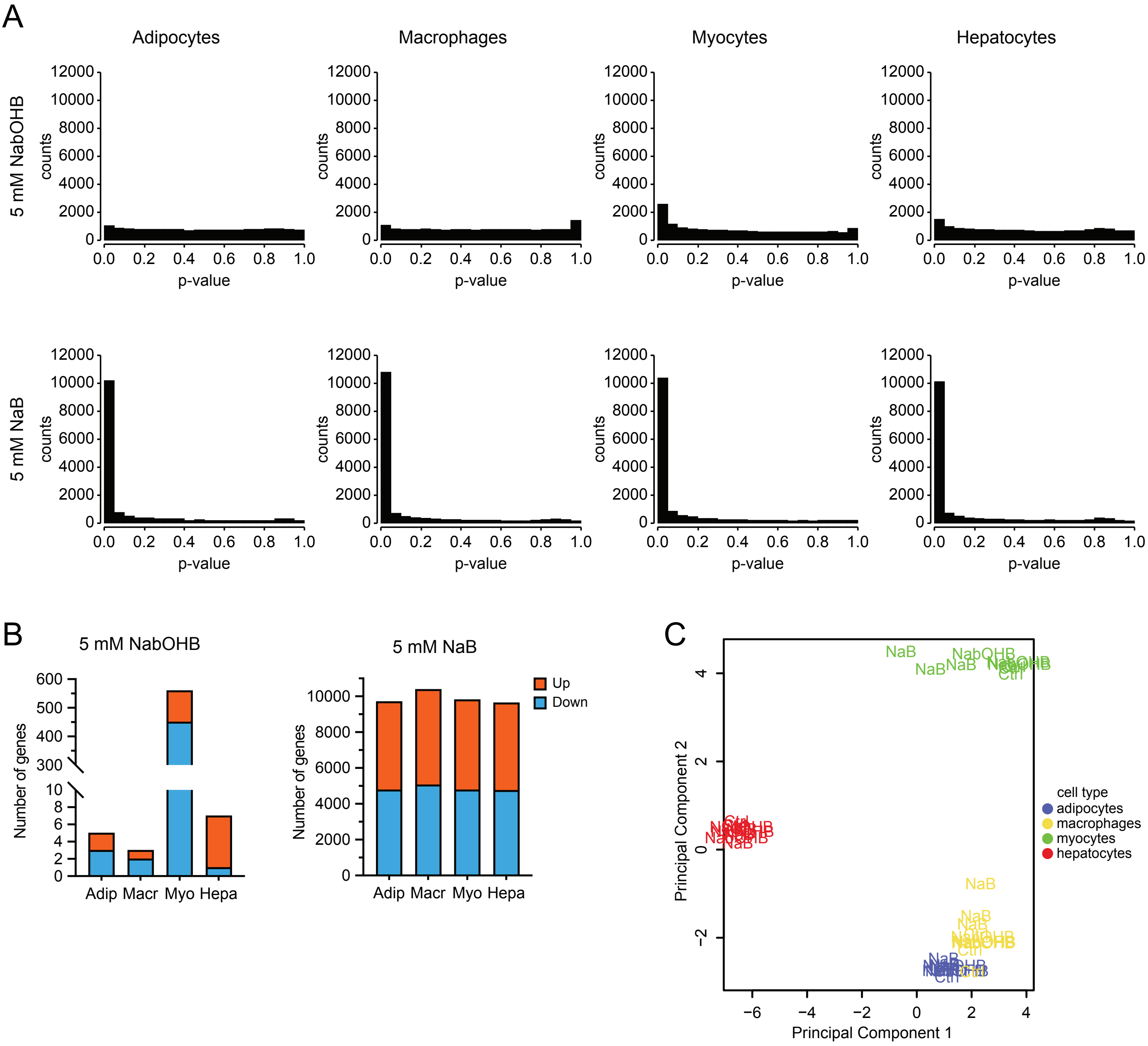


Supplemental figure 2: (A) Expression levels (log2CPM) of monocarboxylate transporters Mct1, Mct2 and Mct7 in relation to Gapdh and Bdh1. (B) Raw p-value histograms for butyrate and βOHB treated cell types. (C) Number of genes significantly altered by treatment with βOHB and butyrate (FDR q < 0.05). (D) Principle component analysis of βOHB and butyrate treated samples.
