## Supplemental tables 1-3 for "Transcriptomic analysis reveals niche gene expression effects of beta-hydroxybutyrate in primary myotubes"

**Supplemental table 1: FDR p<0.001 genes regulated by NabOHB treatment in primary myocytes.**

| Gene name | log2FC | FDR.BH |
|---|---|---|
| *Gm43590* | -4,46 | 3,8E- 03 |
| *Gm48113* | -4,21 | 2,2E- 02 |
| *Clec4a1* | -4,09 | 1,9E- 04 |
| *Gm49339* | -3,93 | 2,2E- 02 |
| *Gm10645* | -3,68 | 2,2E- 02 |
| *Bcl2a1d* | -3,52 | 7,6E- 04 |
| *Tifab* | -3,47 | 2,9E- 04 |
| *Cd300ld* | -3,45 | 5,0E- 04 |
| *Apoc2* | -3,45 | 2,1E- 02 |
| *AW112010* | -3,32 | 1,1E- 02 |
| *Gm29100* | -3,24 | 7,5E- 04 |
| *Ggt5* | -3,08 | 8,9E- 03 |
| *Osm* | -3,06 | 1,7E- 03 |
| *Pf4* | -3,05 | 1,5E- 03 |
| *Gm21188* | -3,03 | 1,3E- 03 |
| *B430306N03Rik* | -2,99 | 5,6E- 03 |
| *Mfng* | -2,94 | 1,4E- 03 |
| *Prdm1* | -2,88 | 3,0E- 03 |
| *Lrrc25* | -2,87 | 8,4E- 04 |
| *Tnfsf8* | -2,86 | 1,1E- 02 |
| *Gpr34* | -2,86 | 2,0E- 03 |
| *Cd86* | -2,83 | 8,8E- 03 |
| *Acod1* | -2,82 | 2,5E- 02 |
| *Tnfsf13* | -2,82 | 9,0E- 03 |
| *H2-Aa* | -2,81 | 8,3E- 03 |
| *3930402G23Rik* | -2,76 | 3,1E- 02 |
| *Rab19* | -2,76 | 2,4E- 02 |
| *Aoah* | -2,75 | 1,6E- 03 |
| *Mcoln2* | -2,74 | 4,5E- 03 |
| *Card9* | -2,74 | 6,1E- 03 |
| *Apol7c* | -2,70 | 4,4E- 03 |
| *H2-Eb1* | -2,67 | 4,6E- 04 |
| *P2ry13* | -2,64 | 2,7E- 03 |
| *Siglece* | -2,62 | 3,4E- 02 |
| *Slfn5* | -2,61 | 2,7E- 02 |
| *Cd74* | -2,60 | 1,5E- 02 |
| *Kcnj10* | -2,59 | 4,6E- 04 |
| *Clec10a* | -2,57 | 2,6E- 02 |
| *Abcg3* | -2,54 | 5,0E- 04 |
| *Ccr2* | -2,54 | 8,8E- 03 |
| *P2ry12* | -2,51 | 3,4E- 03 |
| *Gmfg* | -2,51 | 1,7E- 03 |
| *Ptpro* | -2,51 | 2,5E- 02 |
| *Inpp5d* | -2,50 | 2,8E- 02 |
| *Tpbgl* | -2,50 | 8,9E- 04 |
| *A630001G21Rik* | -2,48 | 4,1E- 03 |
| *Gpr171* | -2,47 | 2,1E- 02 |
| *Gm5431* | -2,46 | 8,0E- 03 |
| *A130077B15Rik* | -2,43 | 6,8E- 03 |
| *Psmb8* | -2,43 | 1,6E- 02 |
| *Themis2* | -2,41 | 9,1E- 03 |
| *Clec7a* | -2,40 | 2,1E- 03 |
| *Gna15* | -2,37 | 4,4E- 02 |
| *Naip6* | -2,36 | 7,3E- 03 |
| *Fcgr2b* | -2,36 | 1,8E- 03 |
| *Srgn* | -2,34 | 4,6E- 04 |
| *Itgax* | -2,34 | 1,6E- 03 |
| *Fcgr4* | -2,33 | 5,6E- 03 |
| *Ms4a4a* | -2,33 | 5,7E- 03 |
| *H2-Ab1* | -2,33 | 3,2E- 02 |
| *Slamf8* | -2,33 | 3,7E- 02 |
| *Cx3cr1* | -2,31 | 8,9E- 04 |
| *Gm11992* | -2,31 | 7,3E- 03 |
| *Lst1* | -2,30 | 1,4E- 02 |
| *Sncaip* | -2,28 | 8,9E- 03 |
| *2210406H18Rik* | -2,27 | 1,1E- 02 |
| *Clec4n* | -2,27 | 3,8E- 03 |
| *Csf2rb* | -2,25 | 1,4E- 02 |
| *Tlr9* | -2,25 | 2,6E- 02 |
| *Icam2* | -2,24 | 3,7E- 02 |
| *Apobec1* | -2,24 | 2,2E- 04 |
| *Batf* | -2,24 | 2,9E- 02 |
| *Tnfrsf11a* | -2,24 | 1,7E- 03 |
| *4933430I17Rik* | -2,23 | 2,3E- 02 |
| *Ms4a14* | -2,23 | 2,0E- 03 |
| *C5ar1* | -2,22 | 1,7E- 03 |
| *Spi1* | -2,21 | 8,8E- 03 |
| *C1qa* | -2,21 | 1,8E- 03 |
| *Drd1* | -2,21 | 1,3E- 02 |
| *Clec4a3* | -2,21 | 1,6E- 02 |
| *Cnr2* | -2,20 | 1,3E- 02 |
| *Fgd2* | -2,19 | 7,3E- 03 |
| *Fcrls* | -2,19 | 9,9E- 04 |
| *Ms4a6c* | -2,18 | 4,6E- 04 |
| *B3gnt7* | -2,15 | 7,6E- 03 |
| *Cd244a* | -2,15 | 4,0E- 02 |
| *Btk* | -2,15 | 6,0E- 04 |
| *Siglec1* | -2,14 | 3,5E- 02 |
| *Plcb2* | -2,14 | 2,1E- 02 |
| *Cysltr1* | -2,13 | 1,7E- 03 |
| *Selplg* | -2,13 | 1,6E- 02 |
| *Lyl1* | -2,12 | 3,7E- 02 |
| *Tlr13* | -2,11 | 1,0E- 03 |
| *Arhgap9* | -2,10 | 7,5E- 04 |
| *Grap2* | -2,10 | 2,2E- 02 |
| *Pik3cg* | -2,10 | 6,8E- 03 |
| *Cd200r2* | -2,10 | 2,2E- 02 |
| *Slc15a3* | -2,09 | 1,6E- 02 |
| *C1qc* | -2,08 | 5,7E- 04 |
| *She* | -2,07 | 3,7E- 02 |
| *Tyrobp* | -2,06 | 7,5E- 04 |
| *Il7r* | -2,06 | 2,3E- 02 |
| *Rps2 -ps13* | -2,05 | 1,6E- 02 |
| *Cybb* | -2,01 | 2,6E- 03 |
| *Fermt3* | -2,01 | 5,7E- 04 |
| *Dock2* | -2,01 | 1,4E- 03 |
| *Ccr1* | -2,00 | 7,2E- 04 |
| *Vav1* | -2,00 | 1,6E- 03 |
| *Ms4a6d* | -1,99 | 9,8E- 04 |
| *Nlrp1b* | -1,99 | 3,3E- 03 |
| *Ptprc* | -1,97 | 4,6E- 03 |
| *C1qb* | -1,97 | 6,3E- 04 |
| *Fcrl1* | -1,95 | 4,3E- 02 |
| *Stap1* | -1,95 | 4,0E- 02 |
| *Tnf* | -1,95 | 1,0E- 02 |
| *Foxj1* | -1,94 | 3,3E- 02 |
| *Grk3* | -1,93 | 5,0E- 03 |
| *Napsa* | -1,92 | 2,2E- 02 |
| *Cd52* | -1,92 | 2,1E- 03 |
| *Irf8* | -1,90 | 1,4E- 03 |
| *Cd33* | -1,90 | 3,8E- 03 |
| *Slc11a1* | -1,90 | 1,4E- 02 |
| *Gm37168* | -1,90 | 2,1E- 02 |
| *Il10ra* | -1,89 | 4,9E- 02 |
| *Gpr65* | -1,89 | 2,5E- 02 |
| *Bcl2a1b* | -1,89 | 1,9E- 03 |
| *Lair1* | -1,88 | 2,5E- 02 |
| *Ebi3* | -1,88 | 3,6E- 02 |
| *Runx3* | -1,88 | 8,1E- 03 |
| *Tal1* | -1,88 | 4,1E- 02 |
| *Csf1r* | -1,88 | 1,3E- 03 |
| *Hcls1* | -1,87 | 2,2E- 03 |
| *Cd300c2* | -1,87 | 1,1E- 02 |
| *Csf2rb2* | -1,86 | 1,0E- 02 |
| *Lpxn* | -1,86 | 4,5E- 03 |

| Gene name | log2FC | FDR.BH | Gene name | log2FC | FDR.BH | Gene name | log2FC | FDR.BH |
|---|---|---|---|---|---|---|---|---|
| *Igsf6* | -1,86 | 1,7E-03 | *Cd68* | -1,69 | 2,5E-03 | *Lyz2* | -1,52 | 9,8E-04 |
| *Stra6l* | -1,86 | 4,8E-02 | *Lilr4b* | -1,68 | 3,7E-02 | *Otulinl* | -1,52 | 4,0E-02 |
| *Tnfaip8l2* | -1,86 | 2,8E-03 | *Nuf2* | -1,68 | 3,9E-03 | *Rassf6* | -1,52 | 4,7E-02 |
| *Cd83* | -1,85 | 1,6E-03 | *Arhgap25* | -1,67 | 6,5E-03 | *BE692007* | -1,51 | 4,3E-02 |
| *Casp1* | -1,85 | 1,0E-02 | *Dpep2* | -1,67 | 1,1E-02 | *Itgam* | -1,51 | 1,3E-02 |
| *I830077J02Rik* | -1,85 | 1,4E-02 | *Tlr1* | -1,66 | 4,5E-03 | *Ncf2* | -1,50 | 1,3E-02 |
| *Bin2* | -1,84 | 3,3E-02 | *Rasgef1b* | -1,66 | 1,2E-02 | *Fgr* | -1,50 | 4,1E-02 |
| *Coro1a* | -1,84 | 8,8E-03 | *C3ar1* | -1,66 | 7,5E-04 | *Il2rg* | -1,50 | 3,8E-03 |
| *Naip2* | -1,84 | 1,8E-02 | *Dok3* | -1,65 | 1,9E-03 | *Tmem273* | -1,50 | 8,8E-03 |
| *Was* | -1,84 | 1,2E-02 | *Rinl* | -1,65 | 4,2E-03 | *Esco2* | -1,48 | 3,0E-02 |
| *Stab1* | -1,83 | 6,8E-03 | *Pld4* | -1,65 | 3,2E-03 | *Tmem106a* | -1,47 | 3,8E-03 |
| *Hk3* | -1,83 | 1,3E-02 | *Cxcl16* | -1,64 | 1,7E-03 | *Lcp2* | -1,47 | 3,5E-02 |
| *Ctss* | -1,83 | 5,7E-04 | *Alox5ap* | -1,63 | 2,1E-02 | *Kif18b* | -1,44 | 2,4E-02 |
| *Ccdc88b* | -1,82 | 4,4E-02 | *Evi2b* | -1,63 | 4,2E-02 | *Ptpn6* | -1,42 | 1,7E-03 |
| *Myo1f* | -1,82 | 8,7E-04 | *Ccl9* | -1,63 | 3,0E-02 | *Clec4d* | -1,40 | 1,4E-02 |
| *Fcgr1* | -1,82 | 2,4E-02 | *Aif1* | -1,63 | 8,8E-03 | *Sla* | -1,39 | 8,7E-03 |
| *Lcp1* | -1,80 | 1,4E-03 | *Pirb* | -1,63 | 5,0E-03 | *Ccl6* | -1,38 | 5,5E-03 |
| *Gpr31c* | -1,80 | 1,3E-02 | *Shtn1* | -1,63 | 9,4E-03 | *Adam8* | -1,38 | 2,2E-04 |
| *Styxl1* | -1,78 | 3,6E-02 | *AI662270* | -1,62 | 1,8E-02 | *Fgl2* | -1,37 | 2,1E-02 |
| *Ikzf1* | -1,78 | 1,1E-02 | *Arhgap30* | -1,62 | 6,6E-03 | *Lgals3bp* | -1,36 | 7,7E-03 |
| *Sirpb1c* | -1,78 | 2,2E-02 | *Ltc4s* | -1,62 | 4,4E-02 | *Cenpa* | -1,36 | 1,4E-02 |
| *Trem2* | -1,77 | 7,5E-04 | *Arhgap45* | -1,62 | 4,6E-04 | *Cd180* | -1,36 | 2,9E-02 |
| *P2ry6* | -1,76 | 2,6E-03 | *Cd200r4* | -1,61 | 7,9E-03 | *Hmgb2* | -1,34 | 1,1E-02 |
| *Marchf1* | -1,75 | 2,1E-02 | *Hck* | -1,60 | 5,0E-04 | *Mmp13* | -1,34 | 2,9E-02 |
| *Plxnc1* | -1,75 | 1,7E-03 | *Tgfbi* | -1,60 | 3,4E-03 | *Fut1* | -1,34 | 4,9E-02 |
| *H2-DMb1* | -1,75 | 2,6E-02 | *Angptl7* | -1,59 | 1,1E-02 | *Icosl* | -1,33 | 3,4E-02 |
| *Entpd1* | -1,75 | 2,3E-03 | *Dock10* | -1,59 | 2,0E-03 | *Arhgap4* | -1,32 | 4,7E-02 |
| *Ms4a7* | -1,74 | 1,7E-03 | *Cd48* | -1,59 | 3,6E-02 | *Cd93* | -1,31 | 2,9E-04 |
| *Snx20* | -1,74 | 1,1E-02 | *Ncf1* | -1,58 | 3,0E-04 | *Cxcl14* | -1,30 | 2,4E-02 |
| *Cd84* | -1,74 | 8,2E-04 | *Neurl3* | -1,57 | 8,7E-04 | *Cfp* | -1,28 | 8,1E-03 |
| *B3gnt8* | -1,73 | 1,9E-02 | *Laptm5* | -1,57 | 5,9E-04 | *Mki67* | -1,27 | 2,1E-02 |
| *Arl11* | -1,73 | 7,7E-03 | *Tlr7* | -1,57 | 8,9E-03 | *Fbxo5* | -1,26 | 2,1E-02 |
| *Gngt2* | -1,73 | 5,6E-03 | *Sash3* | -1,57 | 2,8E-02 | *Ndc80* | -1,26 | 2,5E-02 |
| *Gpr162* | -1,72 | 1,3E-02 | *Cyth4* | -1,56 | 5,7E-04 | *AB124611* | -1,20 | 4,2E-02 |
| *Fcgr3* | -1,72 | 2,2E-04 | *Nrros* | -1,56 | 8,8E-03 | *Elmo1* | -1,19 | 2,7E-02 |
| *Fcer1g* | -1,72 | 1,8E-03 | *Ccl3* | -1,56 | 2,6E-02 | *Cd14* | -1,19 | 2,1E-02 |
| *Pik3ap1* | -1,72 | 1,4E-02 | *Gm14548* | -1,55 | 1,8E-02 | *Top2a* | -1,18 | 1,1E-02 |
| *Il16* | -1,71 | 2,7E-03 | *Mybl2* | -1,55 | 2,5E-02 | *Apobr* | -1,16 | 3,7E-03 |
| *Tbxas1* | -1,71 | 1,7E-02 | *Rasal3* | -1,55 | 3,0E-02 | *Lpcat2* | -1,16 | 2,2E-02 |
| *Mpeg1* | -1,71 | 1,4E-03 | *Plek* | -1,55 | 4,6E-04 | *Tnfsf13os* | -1,16 | 3,2E-02 |
| *Itgb2* | -1,70 | 2,6E-03 | *Hrh1* | -1,54 | 4,4E-02 | *Mtfr2* | -1,16 | 3,0E-02 |
| *Wfdc17* | -1,70 | 1,5E-02 | *Lat2* | -1,53 | 1,7E-02 | *Cep55* | -1,15 | 2,1E-02 |
| *Ptpn7* | -1,70 | 1,7E-03 | *Adgre1* | -1,53 | 2,7E-02 | *Ctla2b* | -1,13 | 2,7E-02 |
| *Nlrc4* | -1,69 | 3,1E-02 | *Hvcn1* | -1,53 | 5,2E-03 | *Duxbl1* | -1,13 | 2,2E-02 |
| *Clec5a* | -1,69 | 1,1E-02 | *Il1rn* | -1,52 | 3,4E-02 | *Cd300a* | -1,11 | 2,5E-02 |

| Gene name | log2FC | FDR.BH | Gene name | log2FC | FDR.BH | Gene name | log2FC | FDR.BH |
|---|---|---|---|---|---|---|---|---|
| *Syk* | -1,10 | 4,3E-04 | *Mylk* | -0,66 | 1,7E-02 | *Coro1c* | -0,43 | 1,1E-03 |
| *Adap1* | -1,10 | 1,5E-02 | *Rgs10* | -0,65 | 1,2E-02 | *Dynap* | -0,42 | 2,7E-02 |
| *Slamf9* | -1,10 | 1,6E-02 | *Dhrs3* | -0,65 | 1,4E-03 | *Arl6ip1* | -0,41 | 1,6E-02 |
| *Gda* | -1,09 | 2,0E-03 | *Pla2g7* | -0,64 | 1,3E-02 | *Rhobtb1* | -0,41 | 2,1E-02 |
| *Irf5* | -1,09 | 1,5E-02 | *Clrn1* | -0,64 | 2,4E-02 | *Ftl1* | -0,41 | 4,4E-02 |
| *Rnasel* | -1,08 | 3,4E-03 | *Rab32* | -0,63 | 4,4E-02 | *Kctd12* | -0,41 | 1,3E-02 |
| *Anpep* | -1,06 | 2,5E-02 | *Lpl* | -0,63 | 3,5E-03 | *Rpa2* | -0,41 | 1,3E-02 |
| *Fli1* | -1,06 | 3,8E-03 | *Igfbp4* | -0,62 | 3,4E-02 | *Arpc1b* | -0,41 | 2,4E-02 |
| *Nusap1* | -1,06 | 5,0E-02 | *Cyp1b1* | -0,62 | 2,6E-02 | *Cnn2* | -0,40 | 1,6E-02 |
| *3010003L21Rik* | -1,05 | 4,3E-02 | *4933421O10Rik* | -0,61 | 2,7E-02 | *Casp6* | -0,40 | 2,6E-02 |
| *Lfng* | -1,02 | 2,2E-02 | *Arhgdib* | -0,61 | 6,2E-03 | *Sri* | -0,40 | 2,6E-02 |
| *Kif22* | -1,01 | 4,9E-02 | *Gzme* | -0,59 | 1,5E-02 | *Dock11* | -0,40 | 2,4E-02 |
| *Sirpa* | -1,00 | 2,6E-06 | *Tcp1l1* | -0,59 | 2,7E-02 | *Anxa8* | -0,39 | 2,0E-02 |
| *Mrc1* | -1,00 | 3,2E-02 | *Rab7b* | -0,58 | 3,0E-02 | *Arl4c* | -0,39 | 4,5E-02 |
| *H2-DMa* | -0,99 | 1,6E-03 | *Ctsc* | -0,58 | 5,0E-04 | *Pnmt* | -0,39 | 4,5E-02 |
| *Pik3r5* | -0,99 | 2,9E-02 | *Lipa* | -0,55 | 2,6E-03 | *Pgm2* | -0,39 | 1,8E-02 |
| *Gpnmb* | -0,98 | 1,1E-03 | *Gpr137b-ps* | -0,55 | 4,1E-03 | *Mymx* | -0,39 | 2,2E-03 |
| *Cd28* | -0,94 | 1,4E-03 | *Cacna2d2* | -0,53 | 1,5E-02 | *Tmsb4x* | -0,38 | 9,1E-03 |
| *Nckap1l* | -0,93 | 5,6E-03 | *Hsd17b11* | -0,53 | 2,4E-02 | *Tpd52* | -0,38 | 3,7E-03 |
| *Cotl1* | -0,92 | 2,2E-04 | *P2rx4* | -0,53 | 1,9E-02 | *Specc1* | -0,38 | 2,7E-02 |
| *Gng2* | -0,92 | 1,7E-02 | *Casp8* | -0,52 | 2,5E-02 | *Wdr75* | -0,38 | 9,9E-03 |
| *Atp6v0d2* | -0,89 | 1,3E-02 | *Stmn1* | -0,52 | 1,4E-03 | *Ctsh* | -0,37 | 5,0E-02 |
| *1700007L15Rik* | -0,89 | 2,9E-02 | *Gm17455* | -0,52 | 2,0E-02 | *Arpc5* | -0,37 | 2,4E-03 |
| *Il6ra* | -0,85 | 2,4E-02 | *Ptgs1* | -0,51 | 1,0E-02 | *Edil3* | -0,37 | 2,1E-03 |
| *Fabp5* | -0,85 | 5,7E-04 | *Tagln2* | -0,50 | 2,8E-03 | *Gpr63* | -0,37 | 3,9E-02 |
| *Hps1* | -0,84 | 1,4E-03 | *Havcr2* | -0,50 | 4,6E-02 | *G6pdx* | -0,37 | 1,7E-02 |
| *Slc16a6* | -0,84 | 2,0E-02 | *Man2b1* | -0,49 | 1,2E-02 | *Wipf1* | -0,36 | 2,4E-02 |
| *Glipr1* | -0,84 | 5,8E-03 | *Ptprj* | -0,48 | 2,5E-02 | *Unc93b1* | -0,36 | 3,7E-02 |
| *Abcb1b* | -0,83 | 2,9E-02 | *Fam20a* | -0,48 | 2,0E-02 | *Gusb* | -0,36 | 8,1E-03 |
| *Jade2* | -0,83 | 8,1E-03 | *Tep1* | -0,47 | 5,6E-03 | *Mylip* | -0,36 | 3,7E-02 |
| *Evi2a* | -0,80 | 1,6E-02 | *Galnt7* | -0,47 | 4,9E-02 | *Hgsnat* | -0,35 | 3,7E-02 |
| *Kcnk2* | -0,80 | 4,9E-03 | *Dab2* | -0,46 | 1,7E-03 | *Mafb* | -0,35 | 8,0E-04 |
| *Cd53* | -0,79 | 1,4E-03 | *Gm2a* | -0,46 | 1,4E-03 | *Prkcd* | -0,35 | 7,6E-03 |
| *Ugt1a7c* | -0,75 | 3,4E-02 | *Dock8* | -0,46 | 1,6E-02 | *Swap70* | -0,34 | 4,2E-02 |
| *Plcg2* | -0,74 | 6,6E-03 | *Emb* | -0,45 | 2,3E-02 | *Ocrl* | -0,34 | 2,6E-02 |
| *H2-D1* | -0,73 | 3,0E-02 | *Tor4a* | -0,45 | 2,4E-02 | *Ctsb* | -0,34 | 3,4E-02 |
| *Arhgap19* | -0,71 | 7,5E-03 | *Hexb* | -0,45 | 1,9E-02 | *Slc7a8* | -0,33 | 4,2E-03 |
| *Abca1* | -0,70 | 7,3E-03 | *Il10rb* | -0,45 | 2,6E-03 | *Sema4c* | -0,33 | 9,0E-03 |
| *9630028I04Rik* | -0,70 | 4,1E-02 | *Col10a1* | -0,45 | 4,7E-02 | *Sptlc2* | -0,33 | 1,3E-02 |
| *Fnip2* | -0,68 | 4,3E-03 | *Hmga2* | -0,44 | 9,4E-03 | *Cdk4* | -0,33 | 6,0E-04 |
| *Fignl1* | -0,68 | 2,6E-02 | *Kcnn4* | -0,44 | 4,5E-02 | *Bri3bp* | -0,32 | 4,3E-02 |
| *Vrk1* | -0,67 | 4,4E-02 | *B2m* | -0,44 | 2,2E-02 | *Tpbg* | -0,32 | 2,2E-02 |
| *Dkk2* | -0,67 | 2,9E-04 | *Atr* | -0,43 | 2,7E-02 | *Actb* | -0,32 | 1,6E-02 |
| *Dcx* | -0,66 | 4,6E-04 | *Poglut3* | -0,43 | 2,4E-02 | *Ptk2b* | -0,31 | 1,6E-02 |
| *Aqp1* | -0,66 | 3,6E-02 | *Srsf3* | -0,43 | 7,5E-04 | *Utp20* | -0,31 | 3,0E-02 |

| Gene name | log2FC | FDR.BH | Gene name | log2FC | FDR.BH | Gene name | log2FC | FDR.BH |
|---|---|---|---|---|---|---|---|---|
| *Pip4k2a* | -0,31 | 1,7E-02 | *Mta2* | -0,15 | 3,0E-02 | *Atg9a* | 0,25 | 2,1E-02 |
| *Dtx4* | -0,30 | 2,6E-02 | *Dnm2* | 0,12 | 4,3E-02 | *Arhgap44* | 0,25 | 2,4E-02 |
| *Nt5m* | -0,30 | 3,0E-02 | *Larp1* | 0,12 | 4,9E-02 | *Hrc* | 0,25 | 3,6E-02 |
| *Manf* | -0,30 | 2,7E-02 | *Cs* | 0,14 | 2,2E-02 | *Ciao2b* | 0,25 | 2,4E-02 |
| *Anxa1* | -0,29 | 5,7E-03 | *Carhsp1* | 0,14 | 2,7E-02 | *Hs6st1* | 0,26 | 4,9E-03 |
| *Iqgap1* | -0,29 | 2,7E-02 | *Mdh2* | 0,15 | 3,0E-02 | *Fut11* | 0,26 | 4,2E-02 |
| *Mgp* | -0,29 | 4,9E-02 | *Fxr2* | 0,16 | 4,1E-02 | *Cyc1* | 0,26 | 3,7E-02 |
| *Sgpl1* | -0,29 | 8,2E-03 | *Npepps* | 0,16 | 1,6E-02 | *Sema3d* | 0,26 | 3,9E-02 |
| *Zfp36l2* | -0,29 | 4,1E-02 | *Inppl1* | 0,17 | 2,7E-02 | *Glce* | 0,26 | 4,9E-02 |
| *Rab8b* | -0,28 | 1,6E-02 | *Atmin* | 0,18 | 4,3E-02 | *Rai2* | 0,27 | 1,1E-02 |
| *Itga6* | -0,28 | 9,0E-03 | *Emc1* | 0,18 | 5,0E-02 | *Stbd1* | 0,27 | 1,4E-02 |
| *Actg1* | -0,28 | 4,9E-02 | *Adgre5* | 0,18 | 3,4E-02 | *Mmp15* | 0,27 | 8,8E-03 |
| *Rap2a* | -0,28 | 2,4E-02 | *Aco2* | 0,18 | 2,1E-02 | *Zfp365* | 0,28 | 1,4E-02 |
| *Cers2* | -0,28 | 4,2E-02 | *Stip1* | 0,18 | 2,5E-02 | *Atn1* | 0,28 | 2,5E-02 |
| *Cndp2* | -0,28 | 3,4E-02 | *Ssbp3* | 0,19 | 2,2E-02 | *Slc38a2* | 0,28 | 7,5E-04 |
| *Hmgcr* | -0,28 | 1,9E-02 | *Prcc* | 0,20 | 2,4E-02 | *Iba57* | 0,28 | 4,9E-02 |
| *Txn1* | -0,28 | 1,6E-02 | *Ube2j2* | 0,20 | 3,9E-02 | *Fgfbp1* | 0,28 | 3,1E-02 |
| *Shc4* | -0,27 | 3,7E-02 | *Aldoa* | -0,73 | 6,5E-01 | *Foxo6* | 0,29 | 2,0E-02 |
| *Vwa5a* | -0,27 | 1,1E-02 | *Zswim8* | 0,21 | 3,5E-02 | *Trim3* | 0,29 | 8,9E-03 |
| *Hexa* | -0,27 | 7,3E-03 | *Irf2bp1* | 0,21 | 2,5E-02 | *Gpatch8* | 0,29 | 3,9E-02 |
| *Grn* | -0,27 | 1,3E-02 | *Ppm1b* | 0,21 | 6,1E-03 | *Pnmal2* | 0,29 | 2,2E-03 |
| *Abhd12* | -0,27 | 1,4E-02 | *Ankrd40* | 0,21 | 4,8E-02 | *Adamtsl4* | 0,29 | 3,9E-02 |
| *Psap* | -0,27 | 7,3E-03 | *Fam53a* | 0,21 | 3,2E-02 | *Psg28* | 0,29 | 3,3E-03 |
| *Gnpda1* | -0,26 | 3,3E-02 | *Kcmf1* | 0,21 | 2,4E-02 | *Mtres1* | 0,30 | 2,1E-03 |
| *Lasp1* | -0,26 | 3,8E-02 | *Opa3* | 0,21 | 1,4E-02 | *Fhl1* | 0,30 | 3,4E-02 |
| *Cfl1* | -0,26 | 2,9E-02 | *Rnf126* | 0,21 | 2,4E-02 | *Sall2* | 0,31 | 4,9E-02 |
| *Elmsan1* | -0,26 | 8,5E-03 | *Ints9* | 0,22 | 1,6E-02 | *Nfic* | 0,31 | 1,1E-02 |
| *Rai14* | -0,25 | 1,7E-02 | *Lrrc20* | 0,22 | 1,6E-02 | *Sema6c* | 0,31 | 4,2E-02 |
| *Tmod3* | -0,25 | 1,8E-02 | *Mtss2* | 0,22 | 3,5E-02 | *Hspa2* | 0,31 | 1,7E-02 |
| *Hnrnpa1* | -0,25 | 2,1E-03 | *Ssx2ip* | 0,22 | 9,0E-03 | *6430548M08Rik* | 0,32 | 2,2E-03 |
| *Marcks* | -0,25 | 2,6E-03 | *L3mbtl3* | 0,22 | 3,7E-02 | *Mafa* | 0,34 | 8,3E-03 |
| *Ralb* | -0,25 | 3,9E-02 | *Stom* | 0,22 | 1,7E-02 | *Traf3ip3* | 0,34 | 3,2E-02 |
| *Gadd45a* | -0,24 | 1,3E-02 | *Dcaf6* | 0,23 | 2,8E-02 | *Fbxl16* | 0,34 | 8,1E-03 |
| *Rap2b* | -0,24 | 1,6E-02 | *Pik3c2b* | 0,23 | 3,2E-02 | *Hivep1* | 0,35 | 2,2E-02 |
| *Snx2* | -0,24 | 3,8E-02 | *Coq9* | 0,23 | 2,6E-02 | *Cstf2* | 0,36 | 2,1E-03 |
| *Serp1* | -0,23 | 3,7E-02 | *Map1b* | 0,23 | 3,9E-02 | *Fgf11* | 0,36 | 2,0E-03 |
| *Lgmn* | -0,23 | 3,7E-02 | *Tbx15* | 0,23 | 2,1E-02 | *Epb41l3* | 0,37 | 2,6E-02 |
| *Dnajc3* | -0,21 | 3,8E-02 | *Usp31* | 0,24 | 2,9E-02 | *E2f2* | 0,38 | 4,5E-03 |
| *Calm2* | -0,21 | 3,8E-02 | *Ttpal* | 0,24 | 4,6E-02 | *Satb1* | 0,38 | 1,5E-02 |
| *Gnai3* | -0,21 | 4,0E-02 | *Cc2d1a* | 0,24 | 1,6E-02 | *Otub2* | 0,38 | 6,1E-03 |
| *Odc1* | -0,21 | 1,1E-02 | *Lrrn1* | 0,24 | 8,8E-03 | *Efna2* | 0,39 | 4,7E-02 |
| *Actr2* | -0,19 | 3,0E-02 | *Agap3* | 0,24 | 1,6E-03 | *Ckmt2* | 0,41 | 2,0E-02 |
| *Man1c1* | -0,18 | 3,8E-02 | *Gab1* | 0,24 | 2,8E-02 | *Ache* | 0,44 | 2,6E-03 |
| *Rasa3* | -0,18 | 3,6E-02 | *Zfp322a* | 0,25 | 5,0E-02 | *Per1* | 0,47 | 3,0E-02 |
| *Eif4a1* | -0,18 | 2,4E-02 | *Ubald1* | 0,25 | 2,9E-02 | *Nfatc2* | 0,51 | 1,6E-03 |

| **Gene name** | **log2FC** | **FDR.BH** |
|---|---|---|
| *Zscan2* | 0,60 | 1,1E-02 |
| *1700013G24Rik* | 0,60 | 3,1E-02 |
| *Ankrd34a* | 0,60 | 3,2E-02 |
| *Kcna7* | 0,64 | 1,7E-03 |
| *Gm49540* | 0,70 | 3,1E-02 |
| *Rgs4* | 0,71 | 3,4E-02 |
| *Stc1* | 0,80 | 3,1E-02 |
| *Gngt1* | 0,84 | 3,5E-02 |
| *Gm50064* | 0,87 | 4,1E-02 |
| *Myh13* | 1,01 | 3,8E-02 |
| *Xlr3b* | 1,44 | 1,5E-02 |
| *Fam131c* | 1,47 | 1,6E-02 |
| *Gm16675* | 1,77 | 2,6E-02 |
| *Gm20554* | 1,84 | 3,5E-02 |
| *A230056P14Rik* | 1,89 | 4,2E-02 |
| *Gria4* | 2,37 | 3,8E-03 |
| *Zdhhc23* | 2,40 | 1,1E-02 |
| *Sowaha* | 2,46 | 1,9E-02 |
| *Rnf43* | 2,57 | 1,7E-02 |
| *Atp7b* | 2,79 | 3,7E-02 |

**Supplemental table 2: GSEA positively enriched pathways by NabOHB treatment in primary myocytes.**

| NAME | NES | FDR q-val |
|---|---|---|
| MMU00020.CITRATE.CYCLE..TCA.CYCLE..KEGG | 2.35 | 0.000 |
| MMU04710.CIRCADIAN.RHYTHM.KEGG | 2.08 | 0.002 |
| MMU01200.CARBON.METABOLISM.KEGG | 1.92 | 0.012 |
| MMU00190.OXIDATIVE.PHOSPHORYLATION.KEGG | 1.91 | 0.010 |
| MMU04260.CARDIAC.MUSCLE.CONTRACTION.KEGG | 1.91 | 0.009 |
| MMU01210.2.OXOCARBOXYLIC.ACID.METABOLISM.KEGG | 1.88 | 0.010 |
| MMU04714.THERMOGENESIS.KEGG | 1.84 | 0.015 |
| MMU01230.BIOSYNTHESIS.OF.AMINO.ACIDS.KEGG | 1.82 | 0.015 |
| MMU00270.CYSTEINE.AND.METHIONINE.METABOLISM.KEGG | 1.81 | 0.016 |
| MMU03015.MRNA.SURVEILLANCE.PATHWAY.KEGG | 1.77 | 0.020 |
| MMU00051.FRUCTOSE.AND.MANNOSE.METABOLISM.KEGG | 1.74 | 0.023 |
| MMU04152.AMPK.SIGNALING.PATHWAY.KEGG | 1.73 | 0.022 |
| MMU05012.PARKINSON.DISEASE.KEGG | 1.71 | 0.025 |
| MMU04922.GLUCAGON.SIGNALING.PATHWAY.KEGG | 1.68 | 0.031 |
| MMU00630.GLYOXYLATE.AND.DICARBOXYLATE.METABOLISM.KEGG | 1.67 | 0.032 |
| MMU04932.NON.ALCOHOLIC.FATTY.LIVER.DISEASE..NAFLD..KEGG | 1.63 | 0.041 |
| MMU05016.HUNTINGTON.DISEASE.KEGG | 1.57 | 0.060 |
| MMU00640.PROPANOATE.METABOLISM.KEGG | 1.56 | 0.066 |
| MMU04136.AUTOPHAGY...OTHER.KEGG | 1.54 | 0.068 |
| MMU00010.GLYCOLYSIS...GLUCONEOGENESIS.KEGG | 1.50 | 0.092 |
| MMU05010.ALZHEIMER.DISEASE.KEGG | 1.50 | 0.089 |
| MMU00620.PYRUVATE.METABOLISM.KEGG | 1.48 | 0.090 |
| MMU04120.UBIQUITIN.MEDIATED.PROTEOLYSIS.KEGG | 1.48 | 0.089 |
| MMU00534.GLYCOSAMINOGLYCAN.SYNTHESIS.HEPARAN.SULFATE.KEGG | 1.47 | 0.092 |
| MMU04910.INSULIN.SIGNALING.PATHWAY.KEGG | 1.45 | 0.096 |

**Supplemental table 3: GSEA negatively enriched pathways by NabOHB treatment in primary myocytes.**

| NAME | NES | FDR q-val |
|---|---|---|
| MMU05150.STAPHYLOCOCCUS.AUREUS.INFECTION.KEGG | -2.70 | 0.000 |
| MMU04061.VIRAL.PROTEIN.INTERACTION.WITH. CYTOKINE.RECEPTOR.KEGG | -2.50 | 0.000 |
| MMU04145.PHAGOSOME.KEGG | -2.49 | 0.000 |
| MMU04142.LYSOSOME.KEGG | -2.35 | 0.000 |
| MMU04640.HEMATOPOIETIC.CELL.LINEAGE.KEGG | -2.35 | 0.000 |
| MMU05140.LEISHMANIASIS.KEGG | -2.34 | 0.000 |
| MMU05323.RHEUMATOID.ARTHRITIS.KEGG | -2.29 | 0.000 |
| MMU04380.OSTEOCLAST.DIFFERENTIATION.KEGG | -2.27 | 0.000 |
| MMU05332.GRAFT.VERSUS.HOST.DISEASE.KEGG | -2.26 | 0.000 |
| MMU04062.CHEMOKINE.SIGNALING.PATHWAY.KEGG | -2.26 | 0.000 |
| MMU05152.TUBERCULOSIS.KEGG | -2.24 | 0.000 |
| MMU04940.TYPE.I.DIABETES.MELLITUS.KEGG | -2.23 | 0.000 |
| MMU04514.CELL.ADHESION.MOLECULES..CAMS..KEGG | -2.22 | 0.000 |
| MMU04060.CYTOKINE.CYTOKINE.RECEPTOR.INTERACTION.KEGG | -2.21 | 0.000 |
| MMU00531.GLYCOSAMINOGLYCAN.DEGRADATION.KEGG | -2.19 | 0.000 |
| MMU05416.VIRAL.MYOCARDITIS.KEGG | -2.17 | 0.000 |
| MMU04672.INTESTINAL.IMMUNE.NETWORK.FOR.IGA.PRODUCTION.KEGG | -2.16 | 0.000 |
| MMU05330.ALLOGRAFT.REJECTION.KEGG | -2.16 | 0.000 |
| MMU04666.FC.GAMMA.R.MEDIATED.PHAGOCYTOSIS.KEGG | -2.15 | 0.000 |
| MMU05133.PERTUSSIS.KEGG | -2.14 | 0.000 |
| MMU04670.LEUKOCYTE.TRANSENDOTHELIAL.MIGRATION.KEGG | -2.14 | 0.000 |
| MMU05320.AUTOIMMUNE.THYROID.DISEASE.KEGG | -2.14 | 0.000 |
| MMU04612.ANTIGEN.PROCESSING.AND.PRESENTATION.KEGG | -2.10 | 0.000 |
| MMU05322.SYSTEMIC.LUPUS.ERYTHEMATOSUS.KEGG | -2.07 | 0.000 |
| MMU05310.ASTHMA.KEGG | -2.02 | 0.000 |
| MMU04650.NATURAL.KILLER.CELL.MEDIATED.CYTOTOXICITY.KEGG | -1.99 | 0.000 |
| MMU05321.INFLAMMATORY.BOWEL.DISEASE..IBD..KEGG | -1.95 | 0.001 |
| MMU04662.B.CELL.RECEPTOR.SIGNALING.PATHWAY.KEGG | -1.94 | 0.001 |
| MMU04064.NF.KAPPA.B.SIGNALING.PATHWAY.KEGG - | 1.92 | 0.001 |
| MMU04611.PLATELET.ACTIVATION.KEGG | -1.91 | 0.001 |
| MMU03030.DNA.REPLICATION.KEGG | -1.89 | 0.002 |
| MMU03010.RIBOSOME.KEGG | -1.88 | 0.002 |
| MMU04210.APOPTOSIS.KEGG | -1.87 | 0.002 |
| MMU00100.STEROID.BIOSYNTHESIS.KEGG | -1.87 | 0.002 |

| | | |
|---|---|---|
| MMU05144.MALARIA.KEGG | -1.85 | 0.003 |
| MMU04621.NOD.LIKE.RECEPTOR.SIGNALING.PATHWAY.KEGG | -1.82 | 0.004 |
| MMU05169.EPSTEIN.BARR.VIRUS.INFECTION.KEGG | -1.82 | 0.004 |
| MMU05142.CHAGAS.DISEASE..AMERICAN.TRYPANOSOMIASIS..KEGG | -1.81 | 0.004 |
| MMU05164.INFLUENZA.A.KEGG | -1.79 | 0.006 |
| MMU00511.OTHER.GLYCAN.DEGRADATION.KEGG | -1.77 | 0.006 |
| MMU05202.TRANSCRIPTIONAL.MISREGULATION.IN.CANCER.KEGG | -1.75 | 0.008 |
| MMU05418.FLUID.SHEAR.STRESS.AND.ATHEROSCLEROSIS.KEGG | -1.73 | 0.010 |
| MMU00480.GLUTATHIONE.METABOLISM.KEGG | -1.73 | 0.011 |
| MMU00603.GLYCOSPHINGOLIPID.BIOSYNTHESIS.GLOBO.AND.ISOGLOBO.SERIES.KEGG | -1.72 | 0.011 |
| MMU04115.P53.SIGNALING.PATHWAY.KEGG | -1.72 | 0.011 |
| MMU04110.CELL.CYCLE.KEGG | -1.71 | 0.012 |
| MMU05163.HUMAN.CYTOMEGALOVIRUS.INFECTION.KEGG | -1.71 | 0.012 |
| MMU04625.C.TYPE.LECTIN.RECEPTOR.SIGNALING.PATHWAY.KEGG | -1.70 | 0.013 |
| MMU05170.HUMAN.IMMUNODEFICIENCY.VIRUS.1.INFECTION.KEGG | -1.69 | 0.014 |
| MMU05143.AFRICAN.TRYPANOSOMIASIS.KEGG | -1.68 | 0.015 |
| MMU05167.KAPOSI.SARCOMA.ASSOCIATED.HERPESVIRUS.INFECTION.KEGG | -1.67 | 0.017 |
| MMU05221.ACUTE.MYELOID.LEUKEMIA.KEGG | -1.67 | 0.017 |
| MMU04620.TOLL.LIKE.RECEPTOR.SIGNALING.PATHWAY.KEGG | -1.67 | 0.017 |
| MMU04664.FC.EPSILON.RI.SIGNALING.PATHWAY.KEGG | -1.67 | 0.017 |
| MMU04971.GASTRIC.ACID.SECRETION.KEGG | -1.66 | 0.018 |
| MMU00600.SPHINGOLIPID.METABOLISM.KEGG | -1.65 | 0.019 |
| MMU04810.REGULATION.OF.ACTIN.CYTOSKELETON.KEGG | -1.65 | 0.019 |
| MMU04015.RAP1.SIGNALING.PATHWAY.KEGG | -1.65 | 0.019 |
| MMU05132.SALMONELLA.INFECTION.KEGG | -1.64 | 0.020 |
| MMU03440.HOMOLOGOUS.RECOMBINATION.KEGG | -1.64 | 0.020 |
| MMU05100.BACTERIAL.INVASION.OF.EPITHELIAL.CELLS.KEGG | -1.64 | 0.020 |
| MMU00520.AMINO.SUGAR.AND.NUCLEOTIDE.SUGAR.METABOLISM.KEGG | -1.63 | 0.023 |
| MMU04750.INFLAMMATORY.MEDIATOR.REGULATION.OF.TRP.CHANNELS.KEGG | -1.62 | 0.023 |
| MMU04924.RENIN.SECRETION.KEGG | -1.61 | 0.025 |
| MMU04610.COMPLEMENT.AND.COAGULATION.CASCADES.KEGG | -1.59 | 0.031 |
| MMU03430.MISMATCH.REPAIR.KEGG | -1.56 | 0.040 |
| MMU04740.OLFACTORY.TRANSDUCTION.KEGG | -1.55 | 0.043 |
| MMU04979.CHOLESTEROL.METABOLISM.KEGG | -1.55 | 0.044 |
| MMU00240.PYRIMIDINE.METABOLISM.KEGG | -1.55 | 0.045 |
| MMU04072.PHOSPHOLIPASE.D.SIGNALING.PATHWAY.KEGG | -1.54 | 0.048 |

| | | |
|---|---|---|
| MMU00980.METABOLISM.OF.XENOBIOTICS.BY.CYTOCHROME.P450.KEGG | -1.53 | 0.049 |
| MMU05340.PRIMARY.IMMUNODEFICIENCY.KEGG | -1.53 | 0.049 |
| MMU04658.TH1.AND.TH2.CELL.DIFFERENTIATION.KEGG | -1.53 | 0.050 |
| MMU05145.TOXOPLASMOSIS.KEGG | -1.53 | 0.050 |
| MMU04970.SALIVARY.SECRETION.KEGG | -1.52 | 0.050 |
| MMU05166.HUMAN.T.CELL.LEUKEMIA.VIRUS.1.INFECTION.KEGG | -1.51 | 0.056 |
| MMU04933.AGE.RAGE.SIGNALING.PATHWAY.IN.DIABETIC.COMPLICATIONS.KEGG | -1.51 | 0.056 |
| MMU04540.GAP.JUNCTION.KEGG | -1.50 | 0.059 |
| MMU00340.HISTIDINE.METABOLISM.KEGG | -1.50 | 0.059 |
| MMU04071.SPHINGOLIPID.SIGNALING.PATHWAY.KEGG | -1.50 | 0.058 |
| MMU00533.GLYCOSAMINOGLYCAN.BIOSYNTHESIS...KERATAN.SULFATE.KEGG | -1.49 | 0.061 |
| MMU04510.FOCAL.ADHESION.KEGG | -1.48 | 0.068 |
| MMU00590.ARACHIDONIC.ACID.METABOLISM.KEGG | -1.48 | 0.068 |
| MMU05014.AMYOTROPHIC.LATERAL.SCLEROSIS..ALS..KEGG | -1.47 | 0.070 |
| MMU04218.CELLULAR.SENESCENCE.KEGG | -1.47 | 0.071 |
| MMU04914.PROGESTERONE.MEDIATED.OOCYTE.MATURATION.KEGG | -1.45 | 0.081 |
| MMU05200.PATHWAYS.IN.CANCER.KEGG | -1.45 | 0.081 |
| MMU00513.VARIOUS.TYPES.OF.N.GLYCAN.BIOSYNTHESIS.KEGG | -1.45 | 0.083 |
| MMU04973.CARBOHYDRATE.DIGESTION.AND.ABSORPTION.KEGG | -1.44 | 0.085 |
| MMU05146.AMOEBIASIS.KEGG | -1.44 | 0.087 |
| MMU05134.LEGIONELLOSIS.KEGG | -1.44 | 0.086 |
| MMU05135.YERSINIA.INFECTION.KEGG | -1.43 | 0.088 |
| MMU00053.ASCORBATE.AND.ALDARATE.METABOLISM.KEGG | -1.43 | 0.089 |
| MMU05032.MORPHINE.ADDICTION.KEGG | -1.43 | 0.091 |
| MMU00514.OTHER.TYPES.OF.O.GLYCAN.BIOSYNTHESIS.KEGG | -1.42 | 0.093 |
| MMU04520.ADHERENS.JUNCTION.KEGG | -1.42 | 0.093 |
